# TMO: ASYMMETRIC CROSS-MODAL ATTENTION FOR LEARNING CELL-STATE-DEPENDENT REGULATORY LAGS FROM SINGLE-CELL MULTIOMIC DATA

**DOI:** 10.64898/2026.06.08.730880

**Authors:** Phabel A. Lopez-Delgado, Mirna M. Delgado-Carlo

**Affiliations:** Center for Genomic Sciences, National Autonomous University of Mexico (UNAM), Morelos, Mexico; Universidad Nacional Autónoma de México, Mexico City, Mexico

**Keywords:** single-cell multi-omics, chromatin accessibility, gene expression, regulatory lag, asymmetric attention

## Abstract

**Background:** Single-cell multi-omics technologies simultaneously measure chromatin accessibility (ATAC) and gene expression (RNA), providing a unique window into the temporal ordering of regulatory events during differentiation. However, most computational models treat the two modalities symmetrically, ignoring the directional relationship between chromatin and transcription, and existing lag-aware methods estimate a single global lag per gene, failing to capture cell-state-dependent dynamics.

**Methods and Results:** We introduce Temporal Multi-Omics (TMO), a deep learning framework that learns signed, cell-state-conditional regulatory lags (Δ*τ*) using asymmetric cross-modal attention. TMO projects RNA and ATAC into 50 latent components each, tokenises each cell as a sequence of 100 tokens, and uses a two-pass transformer in which a data-driven lag prior – derived from a sliding-window cross-correlation function – directly biases attention asymmetrically. On four independent 10x Multiome datasets (mouse brain, human brain, mouse kidney, human PBMC), the asymmetric model achieves Lag Concordance Scores (LCS) of 0.988–0.999, compared to 0.048–0.108 for an architecturally identical symmetric baseline. A stratified 80/20 held-out experiment confirms that the learned component-lag ordering generalises to unseen cells (held-out LCS 0.85–0.99). Clustered Δ*τ* heatmaps show positive Δ*τ* (ATAC-led priming) in early pseudotime and negative Δ*τ* (RNA-led, activity-dependent regulation) in late pseudotime; the ATAC-RNA correlation heatmap exhibits a U-shaped pattern indicative of developmental decoupling. Components with the most positive Δ*τ* are enriched for chromatin organization and stem cell differentiation (FDR < 0.05), while those with the most negative Δ*τ* are enriched for synaptic signalling and immune activation. Ablating the cell-state information from the lag predictor reduces the LCS and collapses per-component temporal dynamics (KS *p* ≤ 0.039 in all four tissues), proving that TMO’s dynamic lag patterns depend on cell-state conditioning. Independent ChIP-seq validation for four transcription factors (PAX5, Pax6, ASCL1, Hnf4*α*) confirms highly significant separation between target genes and expression-matched background (*p <* 10^−4^ in all cases). Two Multiome Perturb-seq screens provide causal validation: SMARCB1 knockout shows a directional trend (1.5-fold target shift, *p* = 0.056, *n* = 147 perturbed cells), and SMARCE1 knockout reaches statistical significance (*p* = 0.0089, *n* = 3,394 perturbed cells). Gene-level cross-correlation independently validates that the regulatory lag signal is present in the raw data, and TMO further identifies rare, statistically significant biphasic gene programs where the regulatory direction reverses across pseudotime.

**Conclusions:** **T**MO is the first method to make regulatory lag a learnable, cell-state-conditional, and architecturally encoded parameter. It is scalable, interpretable, and open-source, providing a powerful tool for studying regulatory timing in development, disease, and perturbation screens.

**Highlights:**

1. TMO learns signed, cell-state-conditional regulatory lags (Δ*τ*) from paired ATAC+RNA data.
2. Asymmetric cross-modal attention biases the model towards the direction of regulation (ATAC leads RNA or RNA leads ATAC).
3. The asymmetric model achieves Lag Concordance Scores > 0.98, far outperforming a symmetric baseline (< 0.11).
4. Stratified 80/20 held-out experiments across all four tissues show that the learned component-lag ordering transfers to unseen cells (held-out LCS 0.85–0.99).
5. Perturb-seq screening detects perturbation-induced shifts in regulatory timing: SMARCB1 knockout shows a directional trend (*p* = 0.056), and SMARCE1 knockout reaches significance (*p* = 0.0089), demonstrating causal relevance.
6. Independent ChIP-seq validation across four transcription factors and tissues confirms that TMO-derived lags distinguish physically bound genes from expression-matched background (*p <* 10^−4^ in all cases).
7. Removing the cell embedding from the LagMLP causes a substantial drop in LCS and collapses the per-component temporal dynamics, proving that TMO’s cell-state conditioning is essential.
8. TMO is open-source, scalable, and ready for application to any 10x Multiome dataset.
9. Gene-level cross-correlation confirms the lag signal is intrinsic to the data and is 3.7–4.7× more variable than TMO’s denoised component profiles.
10. TMO identifies statistically significant biphasic gene programs whose regulatory direction reverses across pseudotime, a new dynamic inaccessible to existing scalar methods.

**Figure 1: Graphical Abstract:** Figure 1:
Graphical Abstract | TMO – Temporal Multi-Omics.
TMO uses asymmetric cross-modal attention in a two-pass transformer to learn signed, cell-state-conditional regulatory lags (Δ*τ*) between chromatin accessibility (ATAC) and gene expression (RNA) from paired single-cell multi-omic data. A data-driven lag prior derived from a sliding-window cross-correlation function biases ATAC-to-RNA attention, enabling the model to capture ATAC-led priming (positive Δ*τ*, red) early in differentiation and RNA-led, activity-dependent regulation (negative Δ*τ*, blue) late in differentiation. Across four tissues, the asymmetric model achieves Lag Concordance Scores (LCS) of 0.988–0.999, while a symmetric baseline collapses to <0.11. Independent ChIP-seq and Perturb-seq validations confirm that the learned lags reflect genuine transcription factor binding and respond to genetic perturbations. TMO is open-source, scalable, and available as a Scanpy-like Python package (tmopy). Made with BioRender.

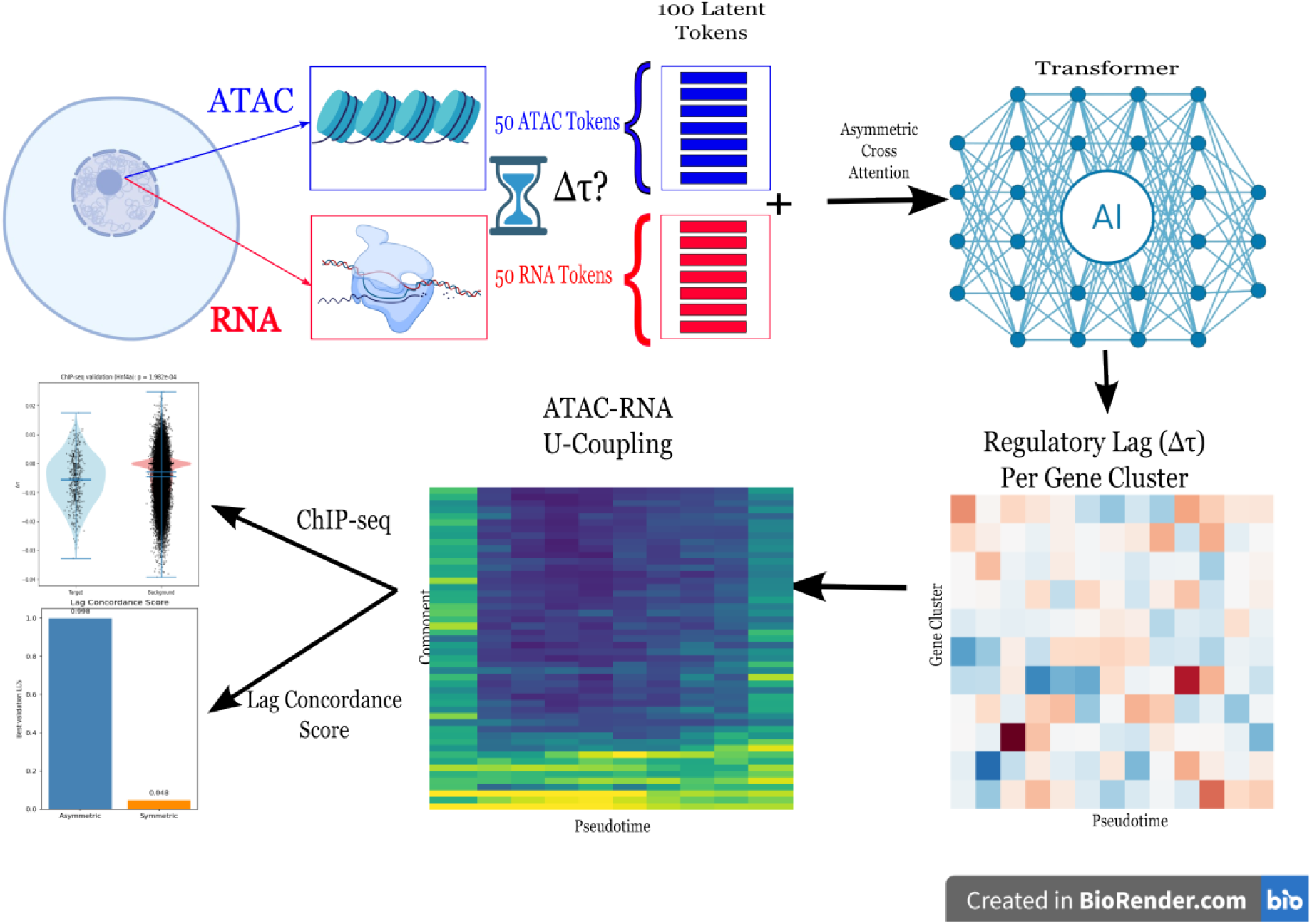

## 2. Introduction

Cell differentiation is a fundamentally temporal process. As a progenitor cell commits to a specific lineage, it traverses a regulatory cascade in which changes to chromatin accessibility precede and enable subsequent changes in gene expression. This epigenetic priming mechanism, in which transcription factors bind to regulatory elements, open chromatin, and license transcription of target genes, has been documented across diverse developmental systems, from haematopoiesis and neurogenesis to kidney organogenesis [1–3]. Yet despite the widespread availability of technologies that simultaneously capture chromatin accessibility and gene expression in the same single cells, the computational tools capable of precisely quantifying *when* and *to what degree* chromatin accessibility leads gene expression, and how this temporal relationship shifts as cells differentiate, remain limited. Addressing this gap is essential for understanding how the epigenome encodes developmental timing, and for predicting how perturbations to chromatin regulators propagate into transcriptional dysregulation in disease.

The development of single-cell multi-omic technologies, in particular the 10x Genomics Multiome platform, has made it routine to obtain paired measurements of chromatin accessibility (via ATAC-seq) and gene expression (via RNA-seq) from thousands of individual cells [1]. These paired datasets carry an untapped temporal signal: because ATAC and RNA are captured from the same cell, differences in their progression along a pseudotime trajectory can directly reveal the regulatory lag between epigenomic priming and transcriptional activation. Pseudotime methods, particularly diffusion pseudotime [4] as implemented in Scanpy [5], provide a principled, root-free ordering of cells along developmental axes, enabling the analysis of gene expression and chromatin dynamics as a continuous function of developmental progression. However, most existing computational frameworks treat ATAC and RNA as interchangeable modalities, fusing them symmetrically without regard for their causal relationship [6, 7]. This symmetry assumption is not merely a simplification, it is biologically incorrect in a directional sense, and it prevents models from learning anything about the regulatory timing that encodes developmental logic.

A small number of methods have begun to address the temporal asynchrony between chromatin and transcription. MultiVelo [8] extends RNA velocity to incorporate chromatin data via a system of ordinary differential equations, estimating a per-gene switch time that reflects the lag between ATAC and RNA dynamics. GrID-Net [9] introduces the concept of “cell-state parallax” and applies Granger causality on DAG-structured trajectories to infer causal locus-gene associations with an explicit lag term. MoFlow [10] proposes a relay velocity framework with locally adaptive, cell-state-dependent kinetics that distinguishes chromatin-dependent from chromatin-independent transcriptional regulation. DELAY [11] uses a convolutional neural network trained on pseudotime-lagged trajectory matrices to infer regulatory relationships, and Velorama [12] applies Granger causal inference on DAG-structured trajectories to identify interaction speeds between transcription factors and their targets. Despite their contributions, these methods share a fundamental limitation: regulatory lag is estimated as a global or trajectory-level statistic per gene or locus-gene pair, a fixed scalar that cannot capture the sign changes or cell-state-specific dynamics that arise as cells transition from progenitor to terminal states. A method that captures ATAC-led priming in early progenitors, lag convergence at commitment points, and RNA-led activity-dependent regulation in terminally differentiated cells, all within a single unified, generalizable framework, does not yet exist.

In parallel, transformer architectures [13] have revolutionized single-cell biology, enabling large-scale pretraining on diverse cell atlases and transfer learning to specific biological questions [14–16]. Models such as scGPT [15] and Geneformer [16] have demonstrated the power of self-attention for capturing gene co-expression relationships and enabling zero-shot perturbation prediction. Recent benchmarks [7] confirm that multimodal transformer architectures outperform conventional integration methods across a range of tasks. However, existing multimodal single-cell transformers concatenate ATAC and RNA tokens and apply standard symmetric self-attention, with no mechanism to represent the directional regulatory relationship between chromatin and transcription [6, 15]. The architecture is temporally agnostic: ATAC tokens attend to RNA tokens and RNA tokens attend to ATAC tokens with equal weight, with no prior encoding the fact that chromatin changes causally precede transcription. This represents a fundamental mismatch between the model’s inductive bias and the biology it is meant to capture.

Here **TMO** (Temporal Multi-Omics) is introduced, a transformer-based framework that addresses this gap through three key innovations. First, TMO learns a *signed, cell-state-conditional regulatory lag* Δ*τ* per latent gene program, using a two-pass forward architecture in which the first pass generates a cell embedding from ATAC tokens alone, and the second pass uses that embedding to parameterize an asymmetric attention bias that explicitly encodes the expected temporal ordering between chromatin and transcription. Second, the lag is informed by a data-driven prior derived from a sliding-window cross-correlation function (CCF) computed directly from the latent representations of paired ATAC and RNA components, providing a signed, pseudotime-resolved ground truth for training without requiring any external annotation or database. Crucially, this lag is not a fixed scalar per gene: it is a continuously varying, cell-state-conditional quantity that can change sign along the trajectory, capturing the shift from chromatin priming in progenitors to activity-dependent regulation in terminal states. Third, TMO introduces a uniform biological interpretation convention via automatic pseudotime orientation correction, ensuring that positive Δ*τ* always denotes ATAC-led priming and negative Δ*τ* denotes RNA-led activity-dependent regulation, enabling direct cross-dataset comparison across tissues and species.

We validate TMO on four independent 10x Multiome datasets spanning mouse brain, human brain, mouse kidney, and human peripheral blood mononuclear cells (PBMCs), demonstrating Lag Concordance Scores of 0.988– 0.999 against a data-driven CCF prior, compared to 0.048–0.108 for an architecturally identical symmetric baseline. The learned lags recapitulate known developmental biology through GO enrichment of lagged gene programs and marker gene profiles. TMO was further validated by using two independent, orthogonal strategies: a perturbation-based causal validation using Multiome Perturb-seq data [17, 18] in which transcription factor knockout shifts the regulatory lag of known target gene programs, and an independent ChIP-seq validation using publicly available binding data from ChIP-Atlas [19] in which genes physically bound by a tissue-specific transcription factor showsystematically different Δ*τ* from expression-matched background genes (two-sided Mann-Whitney *p <* 3.7 × 10^−4^ in all four datasets). Together, these results establish TMO as the first framework to make regulatory lag a learnable, generalizable, and causally validated architectural parameter of a foundation model for single-cell multi-omics.

## 3. Background

### 3.1. The chromatin-to-transcription regulatory lag in differentiation

Gene regulation during differentiation operates through a temporally ordered cascade. Transcription factors bind to enhancers and promoters, recruit chromatin remodelling complexes, and progressively open compacted chromatin before RNA polymerase can be recruited and transcription initiated [2, 3]. This sequential process means that chromatin accessibility at a regulatory locus typically precedes the transcriptional activation of its target gene by a measurable temporal offset, the regulatory lag Δ*τ*. Evidence for this priming mechanism has accumulated across multiple developmental contexts: in haematopoiesis, lineage-determining transcription factors open chromatin at target loci hours to days before gene induction [2]; in neurogenesis, proneural factors such as Pax6 and ASCL1 establish broad chromatin accessibility in progenitors before peak transcription of neuronal identity genes [1]; and in kidney development, nephron progenitor identity is encoded in chromatin long before the transcriptional execution of nephrogenesis [1].

Critically, the sign of Δ*τ* is not fixed throughout differentiation. In early progenitor states, chromatin is the regulatory bottleneck: accessibility must be established before transcription can proceed, yielding positive Δ*τ* (ATAC leads RNA). As cells approach terminal differentiation, chromatin at active loci becomes stably open and the regulatory bottleneck shifts to signal-dependent transcriptional machinery. In this regime, genes can be rapidly induced by extracellular signals without preceding chromatin changes, a pattern characteristic of immediate early genes in neurons, cytokine genes in activated immune cells, and activity-dependent programs more broadly, yielding near-zero or negative Δ*τ* (RNA leads or co-occurs with ATAC). A complete model of chromatin-to-transcription dynamics must therefore represent Δ*τ* as a continuously varying, signed quantity that can change along the differentiation axis, not as a fixed scalar per gene.

### 3.2. Pseudotime inference from diffusion maps

Pseudotime methods assign each cell a continuous value representing its position along a developmental trajectory, enabling the analysis of molecular dynamics as a function of developmental progression rather than real time [4]. Diffusion pseudotime (DPT), introduced by Haghverdi and colleagues [4] and implemented in Scanpy [5], computes a low-dimensional embedding of cells using the diffusion geometry of the data manifold. The first diffusion component is intrinsically monotonic along the principal axis of transcriptional variation and provides a robust pseudotime estimate that does not require a user-specified root cell, making it particularly suitable for unsupervised, multi-dataset analysis. In TMO, pseudotime is derived from the first diffusion component of the RNA data, scaled to [0, 1], with an optional orientation correction step using a known early marker gene to ensure consistent biological interpretation across datasets [5].

### 3.3. Latent semantic indexing for chromatin accessibility

Single-cell ATAC-seq data is inherently sparse and high-dimensional, with tens of thousands of peaks per cell but typically fewer than five percent non-zero entries per peak [2, 3]. Latent semantic indexing (LSI), comprising term frequency-inverse document frequency (TF-IDF) normalisation followed by truncated singular value decomposition (SVD), is the standard approach for dimensionality reduction of scATAC-seq data [2, 3]. The TF-IDF transformation down-weights peaks accessible in many cells, analogously to how common words are down-weighted in document retrieval, reducing the influence of constitutively open housekeeping loci and emphasising cell-type-specific accessibility patterns. TMO applies LSI to a gene-level ATAC matrix constructed by aggregating peak accessibility over proximal gene annotations, producing 50 latent components that capture co-accessible gene programs and that are directly comparable to the 50 RNA PCA components used as the paired modality input.

### 3.4. Cross-correlation functions for lag estimation along pseudotime

The cross-correlation function (CCF) between two time series *x*(*t*) and *y*(*t*) at lag *ℓ* is defined as CCF(*ℓ*) = E[*x*(*t*) · *y*(*t* + *ℓ*)], and the lag at which the CCF is maximised identifies the temporal offset at which *x* most strongly predicts future *y* [11, 12]. Applied to pseudotime-ordered single-cell data, the CCF provides a principled, non-parametric estimate of the regulatory lag between matched ATAC and RNA components without requiring a parametric model of the underlying dynamics. A sliding-window approach, in which the CCF is computed within narrow pseudotime windows rather than across the full trajectory, relaxes the stationarity assumption and allows the lag to vary as a function of developmental state, capturing the sign changes described above. Critically, the sliding-window CCF is signed: a positive peak lag indicates ATAC leads RNA (priming), and a negative peak lag indicates RNA leads ATAC (activity-dependen regulation). In TMO, the CCF surface serves as a biologically grounded, data-driven prior for training the asymmetric attention mechanism, providing supervision without requiring any external annotation, database, or prior knowledge of gene regulatory networks.

### 3.5. Transformer architectures for single-cell omics

Transformer models [13], originally developed for natural language processing, have emerged as the dominant paradigm for large-scale single-cell analysis [14–16]. The self-attention mechanism computes pairwise compatibility scores between all tokens in a sequence, enabling the model to capture long-range dependencies without the inductive biases of convolutional or recurrent architectures. In single-cell biology, genes or latent components serve as tokens, and attention weights have been interpreted as proxies for genegene regulatory relationships [15, 16]. Geneformer [16] demonstrated that a transformer pretrained on 30 million single-cell transcriptomes can predict the effects of transcription factor perturbations in unseen cell types via transfer learning. scGPT [15] extended this paradigm to multi-omic data by concatenating gene expression and chromatin accessibility tokens into a shared sequence and applying generative pretraining. Recent benchmarks confirm that multimodal transformer architectures outperform conventional integration methods across annotation, perturbation prediction, and batch correction tasks [6, 7, 14].

However, a fundamental architectural limitation persists across all existing multimodal single-cell transformers: cross-modal attention is computed symmetrically, allowing ATAC tokens to attend to RNA tokens and RNA tokens to attend to ATAC tokens with equal, undirected weight. This symmetry is architecturally agnostic to the causal relationship between chromatin and transcription, it cannot represent the fact that ATAC changes precede RNA changes in priming regimes, nor that the magnitude and sign of this temporal offset varies with cell state. TMO addresses this gap by introducing an asymmetric cross-modal attention bias parameterized by a learnable, cell-state-conditional lag Δ*τ*, making the direction of time an explicit, trainable component of the model architecture.

### 3.6. Perturbation screens and causal validation in multi-omics

Genome-scale perturbation screens, particularly Perturb-seq [18], have become a powerful tool for mapping causal gene regulatory relationships at single-cell resolution. The recent extension to paired ATAC+RNA measurement, Multiome Perturb-seq [17], enables, for the first time, the direct observation of how transcription factor perturbations alter both chromatin accessibility and gene expression simultaneously in the same cells. This creates a natural experimental design for causal validation of computational lag estimates: if a model has learned genuine regulatory timing, then knocking out a transcription factor should produce a larger shift in the predicted regulatory lag for its known target genes than for background genes. TMO implements this causal validation framework as a core component of its evaluation pipeline, using the one-sided Mann-Whitney U test to assess whether perturbation-induced lag shifts are specifically elevated in target gene programs. Complementary validation using ChIP-seq data from ChIP-Atlas [19], which provides genome-wide maps of physical transcription factor binding across hundreds of cell types, enables an orthogonal, binding-based assessment of whether TMO-derived lags systematically differentiate genes directly regulated by a given transcription factor from expression-matched background genes, providing independent evidence that the learned lags reflect genuine regulatory relationships rather than statistical artefacts of the latent space geometry.

## 4. Methods

This section describes the full mathematical framework, algorithms, and biological interpretation of the Temporal Multi-Omics (TMO) model.

### 4.1. Data preprocessing and gene-level ATAC matrix

Each 10x Multiome dataset provides a filtered feature-barcode matrix (HDF5) and an ATAC peak annotation file (TSV). The H5 file is read with `muon.read_10x_h5()` to obtain a MuData object containing two modalities: `rna` (cells × genes) and `atac` (cells × peaks). The annotation TSV maps each peak to its proximal gene.

Let *P* be the number of peaks, *G* the number of unique genes. Construct a binary peak-gene incidence matrix *M* ∈{0, 1 ^*P ×G*^} where *M*_*p,g*_ = 1 if peak *p* is annotated to gene *g*. The gene-level ATAC accessibility matrix (cells× genes) is computed as:

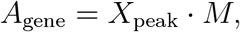

where *X*_peak_ is the cells × peaks count matrix (sparse). *A*_gene_ is stored in `.obsm[‘ATAC_gene’]` as a sparse CSR matrix. Raw RNA counts remain in `.X`.

### 4.2. Pseudotime from first diffusion component and orientation correction

RNA counts are normalised to 10, 000 total counts per cell, log_1_ *p* transformed, and the top 2, 000 highly variable genes are selected (Seurat flavour). PCA is performed with 30 components, followed by a *k*-nearest-neighbour graph (*k* = 15) and diffusion maps (15 components) using `scanpy.tl.diffmap`. The first diffusion component *τ*_0_ is monotonic along the main axis of variation. It is min-max scaled to [0, 1].

The first diffusion component has an arbitrary direction. To enforce biological consistency, an optional early marker gene (e.g., *Pax6* for neurogenesis, *CD34* for haematopoiesis) is provided. Let *x*_*m*_ be the expression vector of the marker. Compute the Pearson correlation *r* = corr(*x*_*m*_, *τ*_0_). If *r <* 0, pseudotime is reversed: *τ* = 1 − *τ*_0_, and a flag `.uns[‘pseudotime_reversed’] = True` is stored. Otherwise *τ* = *τ*_0_ and the flag is `False`. This flag is later used to flip the sign of Δ*τ*, guaranteeing the uniform interpretation:

- Positive Δ*τ* = ATAC leads RNA (chromatin priming)
- Negative Δ*τ* = RNA leads ATAC (activity-dependent regulation)

### 4.3. Latent representations for scalability

To avoid quadratic memory scaling with respect to number of genes or peaks, both modalities are projected into 50 latent components.

- RNA: PCA with 50 components on the log-normalised RNA matrix → *Z*_RNA_ ∈ ℝ^*N ×*50^.
- ATAC: Apply TF-IDF transformation to *A*_gene_:

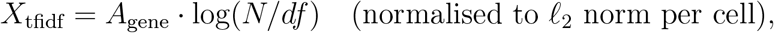

*X*_tfidf_ = *A*_gene_ · log(*N/df*) (normalised to *ℓ*_2_ norm per cell), where *df* is the document frequency (number of cells with non-zero accessibility). Then TruncatedSVD with 50 components → *Z*_ATAC_ ∈ ℝ^*N* ×50^.

Thus each cell is tokenised as a sequence of 100 tokens: first 50 ATAC components, then 50 RNA components. Each token represents a latent gene/accessibility program.

### 4.4. Component annotation (PCA loadings and GO enrichment)

For each RNA component *k* (*k* = 1, …, 50), the top 100 genes by absolute PCA loading are extracted. These gene lists are submitted to Enrichr (`gseapy v1.0`) using the GO_Biological_Process_2023 gene set. The top five GO terms by adjusted *p*-value (Benjamini-Hochberg) are recorded. This provides biological labels for the latent components.

### 4.5. Local cross-correlation function (CCF) for ground-truth lag

For each matched component pair (*i, i*), treat the pseudotime-ordered scores *a*_*i*_(*τ*) (ATAC) and *r*_*i*_(*τ*) (RNA) as locally stationary. A sliding window of half-width *w* = 0.1 is moved across nine centres *τ*_0_ ∈ {0.1, 0.2, …, 0.9}. Inside each window, let *I* = {*t* : |*τ*_*t*_ − *τ*_0_| ≤ *w*}. To handle non-uniform pseudotime spacing, both series are linearly interpolated onto a dense regular grid (200 points), and the cross-correlation is computed using `scipy.signal.correlate` in full mode. The correlation is then mapped back to lag values in pseudotime units, and the peak lag Δ*τ*_*i*_(*τ*_0_) is identified within [− 0.3, 0.3].

The resulting surface Δ*τ*_*i*_(*τ*) is smoothed using Gaussian process regression (Matern-3/2 kernel, bounds: length_scale [10^−2^, 0.5], noise_level [10^−4^, 10^0^]). The final training target for each component is the smoothed value at the last window (*τ* = 0.9):

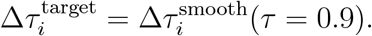

If `.uns[‘pseudotime_reversed’]` is `True`, all 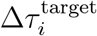 are multiplied by −1.

### 4.6. TMO model architecture (asymmetric cross-modal attention)

The asymmetric model `TMOLatentModelAsymmetric` is a transformer encoder with the following hyperparameters:

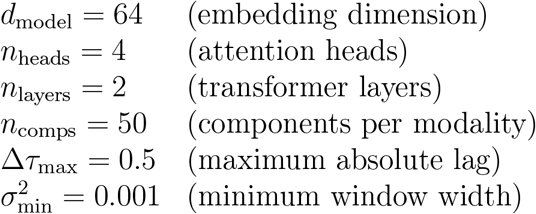

#### Tokenisation

For each cell, tokens are constructed as

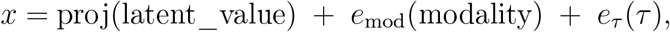

where proj is a linear layer (scalar→*d*_model_) that projects the component’s PCA or LSI score, *e*_mod_ is a learnable modality embedding (0 for RNA, 1 for ATAC), and *e*_*τ*_ (*τ*) is a 2-layer MLP ([128, 64] hidden units, ReLU) that maps the scalar pseudotime *τ* to a *d*_model_-dimensional vector. This design ensures that every token carries cell-state-specific information while also encoding modality and temporal context.

#### Two-pass forward pass

**Pass 1 – cell embedding without bias**. The 100 tokens are processed by the encoder with all cross-attention biases set to zero. After the encoder, the outputs of the 50 ATAC tokens are mean-pooled to obtain a cell embedding 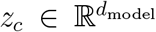. Using ATAC tokens exclusively prevents causal leakage from the RNA stream.

#### Lag and width prediction

For each component index *i*, compute a signed lag 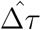 and a squared width 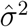 using a shared `LagMLP` and `WidthMLP`. Both take as input the concatenation of *z*_*c*_ and the component embedding *e*_comp_(*i*). The architectures are:

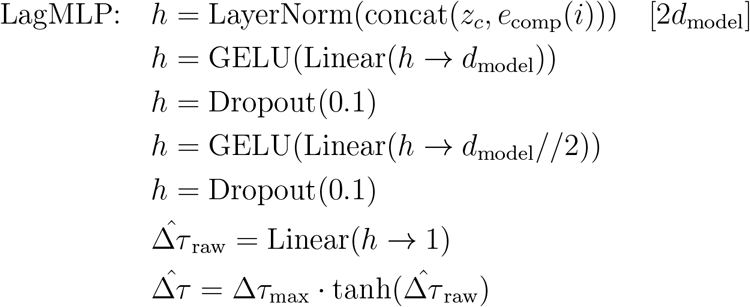

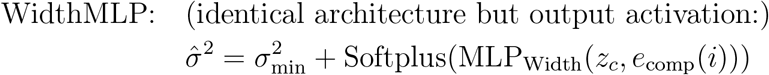

Applied to all 50 indices for both modalities:

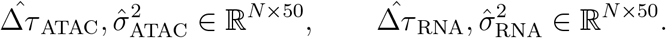

##### Pass 2 – asymmetric biased attention

The same 100 tokens are processed again, but now cross-modal attention biases are active. For an ATAC token *a*_*i*_ at *τ*_*i*_ attending to an RNA token *r*_*j*_ at *τ*_*j*_, the bias is

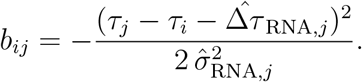

For the reverse direction (RNA token *r*_*j*_ attending to ATAC token *a*_*i*_), the bias is

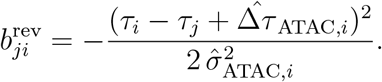

These biases are added to the scaled dot-product attention logits before softmax, making the attention mechanism explicitly asymmetric. Within-modality attention (ATAC-ATAC, RNA-RNA) uses standard unbiased attention.

#### Decoder

After Pass 2, the 50 RNA token outputs are linearly projected to predict the RNA latent components 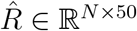.

### 4.7. Training objective

The model is trained on the full dataset (no train/validation split) with a multi-objective loss:

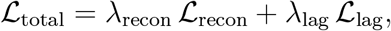

with *λ*_recon_ = 1.0, *λ*_lag_ = 0.1.

Reconstruction loss (MSE):

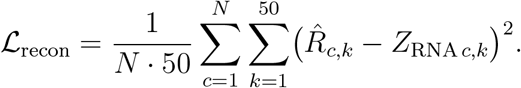

Lag consistency loss: For each ATAC component *i*, compute the mean predicted lag over cells: 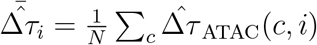. Then

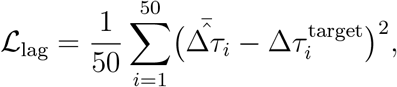

where 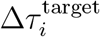 is the smoothed CCF lag at the last pseudotime window (Eq.(4.5)). This loss forces the component-wise ordering of lags to match the CCF-derived order.

Optimiser: Adam (lr = 10^−3^, weight decay = 0.01) with a stepwise learning rate scheduler (gamma = 0.5 at epochs 20 and 40). Training runs for 50 epochs, batch size 32. Every 5 epochs, the Lag Concordance Score (LCS) is computed on the entire dataset (in-set):

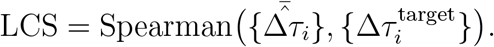

The model checkpoint with the highest LCS is saved as `tmo_asymmetric_best.pt`. The LCS is reported as a diagnostic of how well the model fits the data-derived CCF prior; independent biological validation is provided by ChIP-seq and Perturb-seq experiments.

#### Symmetric baseline

A symmetric baseline is trained with the identical architecture, but the forward pass uses `use_bias = False` (all cross-attention biases are zero) and the lag consistency loss is omitted (*λ*_lag_ = 0). The baseline therefore learns only to reconstruct the RNA latent components and has no mechanism to capture temporal offsets. Its LCS is expected to remain near zero.

### 4.8. Validation metrics

- **Lag Concordance Score (LCS)** measures the Spearman correlation between the model’s predicted per-component lags (averaged over cells) and the CCF targets. It is computed on the full dataset and serves as an in-set measure of internal consistency; high LCS (> 0.9) indicates that the model correctly orders the components by their regulatory lag.
- **ChIP-seq validation**. For a transcription factor, its top-500 bound genes from ChIP-Atlas (peak-to-gene mapping *±*5 kb from TSS) were obtained. Component-level CCF lags are projected to gene-level lags via the PCA loadings: gene_lag = (loadings^⊤^) comp_lag, with the loadings row-normalized to sum to 1. A two-sided Mann-Whitney U test compares the δ*τ* values of target genes with the full set of background genes (all non-target genes, optionally restricted to those within *±*0.5 log-expression units of the target median). A two-sided test was used for ChIP-seq validation, as the direction of δ*τ* is not specified *a priori*. No random subsampling is performed; the test is fully deterministic. Significant p-values indicate that genes physically bound by the TF show a systematically different regulatory lag than background genes. To validate the regulatory lag signal independently of dimensionality reduction, we also computed a gene-by-gene cross-correlation along pseudotime for every gene, using the same sliding-window interpolation procedure applied at the component level (Supplementary Methods).
- **Perturbation-induced lag shift (**δΔ*τ* ). For a Perturb-seq experiment, the asymmetric model is trained on control cells, then applied to perturbed cells. For each ATAC component *i*,

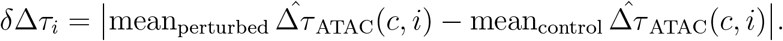

Known target genes are mapped to components via the highest absolute PCA loading. The δΔ*τ* values of target-associated components are compared against all other components using a one-sided Mann-Whitney U test (*α* = 0.05). A one-sided test was used for Perturb-seq validation, reflecting the pre-specified directional hypothesis that perturbation increases target lag shifts. A significant p-value indicates that the perturbation shifts the regulatory lag preferentially for its known targets.
- **Attention Disruption Score (ADS)**. For a TF knockout *k*, let *L*_*k*_ be the set of ATAC peaks containing the TF’s motif. For each gene *g*, the cross-modal attention weight from peak *ℓ* to the RNA token of *g* (averaged over cells) is compared between perturbed and unperturbed conditions:

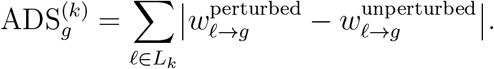

Elevated ADS for known targets relative to background further supports causal relevance.

### 4.9. Biological interpretation of Δτ after uniform sign convention

After applying the automatic sign flip (based on `.uns[‘pseudotime_reversed’]`),

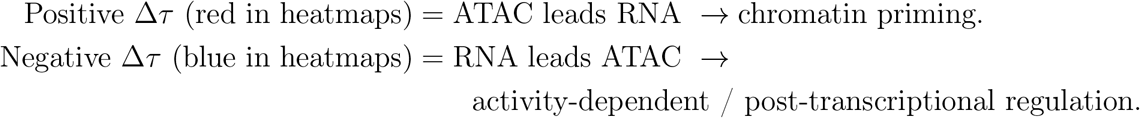

This convention is consistent across all datasets and figures.

### 4.10. ATAC-RNA correlation heatmap

For each RNA component, compute the maximum absolute Pearson correlation with any ATAC component in 10 equal pseudotime bins. The resulting matrix (RNA components × bins) is visualised. A U-shape (high correlation at early and late pseudotime, lower in the middle) indicates developmental decoupling.

### 4.11. Gene program ordering

Components are ranked by Δ*τ* at the last pseudotime window (*τ* = 0.9). The top-10 most positive (ATAC-led) and top-10 most negative (RNA-led) groups are formed. The union of top 100 genes from each group is submitted to GO enrichment. This reveals the biological processes associated with each regulatory mode.

### 4.12. Scalability (windowed sparse attention)

For datasets > 10^4^ cells, the full attention matrix is memory-prohibitive. TMO implements windowed attention that restricts each query token to attend only to keys/values whose pseudotime offset satisfies

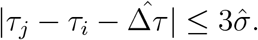

Cells are sorted by pseudotime, partitioned into buckets of size *B*, and attention is computed only between neighbouring buckets. This reduces complexity from *O*(*N* ^2^*L*^2^) to *O*(*NLW*), where *W* is the constant window width.

### 4.13. tmopy: high-level Python API for TMO

To facilitate reproducible and interactive analysis, `tmopy` was developed, a Scanpy-like Python package that provides a unified interface to the TMO framework. The API is organized into three modules:

- `tmopy.pp` (preprocessing): loading 10x Multiome data, constructing gene-level ATAC matrices, computing pseudotime, and optional gene/-peak filtering.
- `tmopy.tl` (tools): component annotation (GO enrichment), training (asymmetric and symmetric), evaluation (Δ*τ* surface, LCS), causal validation (Perturb-seq and ChIP-seq), LCS comparison across models, and gene program ordering.
- `tmopy.pl` (plotting): Δ*τ* heatmaps, ATAC-RNA correlation heatmaps, training curves, violin plots for validation, and marker gene Δ*τ* profiles.

All functions operate on `AnnData` objects and store results in `.obs`, `.obsm`, or `.uns` (e.g., `adata.uns[‘component_annotation’]`, `adata.uns[‘pseudotime_reversed’]`), ensuring full interoperability with Scanpy.

A typical workflow is:

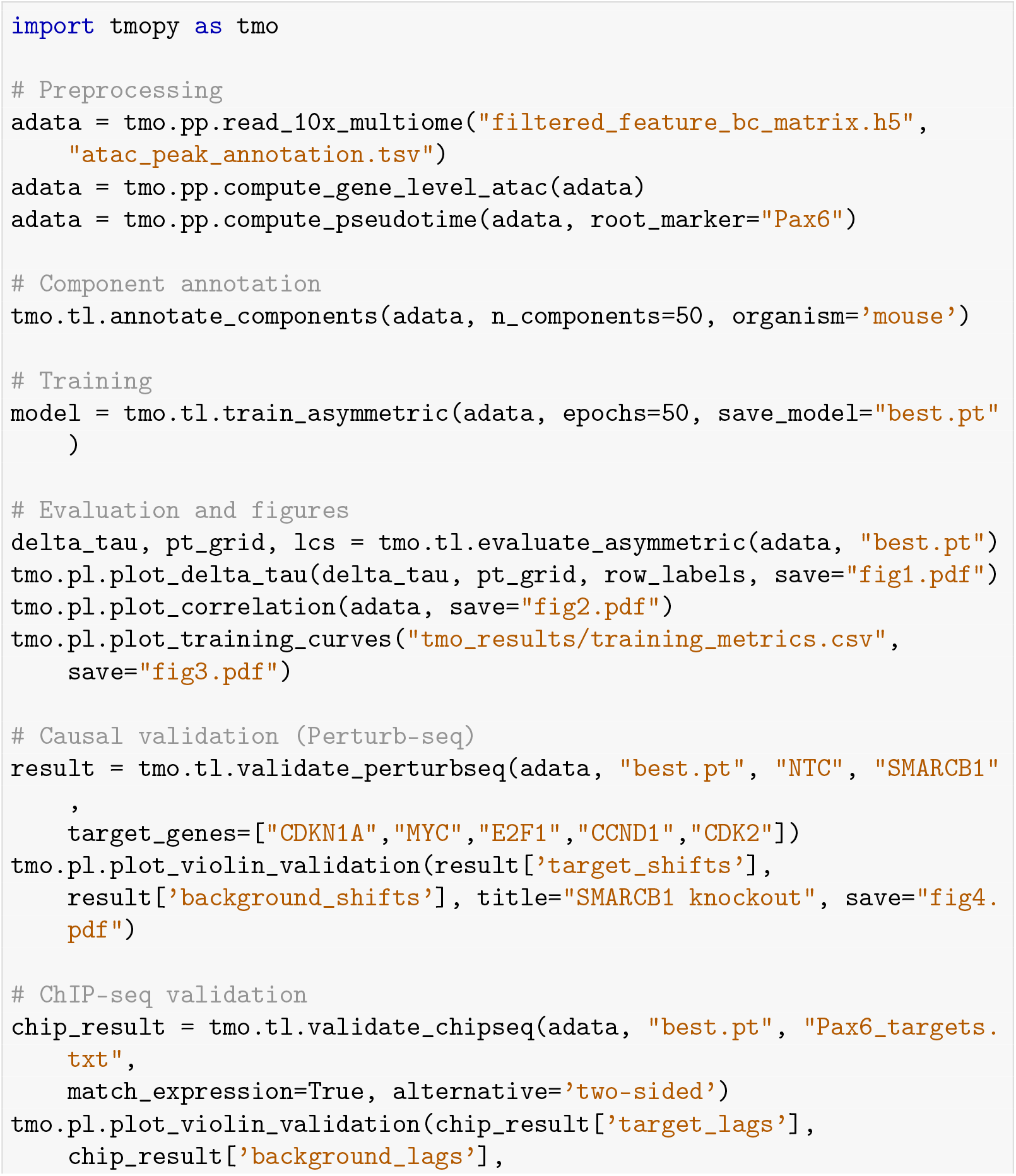
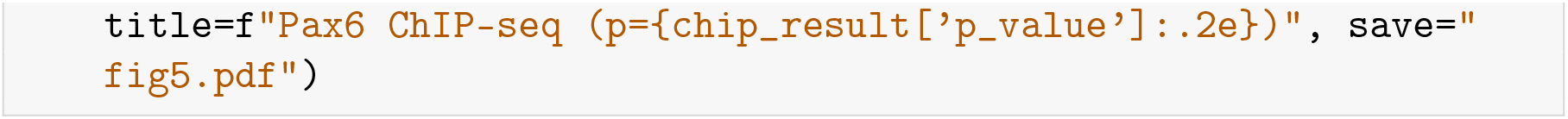

The `tmopy` package is distributed together with the core `tmo` library and installed via “`pip install -e .`“. Detailed documentation is available in the project README and in the docstrings of each function.

## 5. Results

### 5.1. Gene-level ATAC matrix and pseudotime orientation

Mapping peaks to genes produced matrices of size:

- Mouse brain: 4, 881 cells × 31, 036 genes
- Human brain: 3, 332 cells × 30, 496 genes
- Mouse kidney: 5, 287 cells × 24, 712 genes
- Human PBMC: 3, 009 cells × 26, 976 genes

Pseudotime was reversed for mouse brain (using *Pax6*) and mouse kidney (using *Pax2*), while human brain and PBMC were kept in the original orientation. The reversal flag was saved and later used to flip Δ*τ* signs, guaranteeing the uniform biological interpretation.

### 5.2. Asymmetric attention is necessary for learning regulatory lags

The asymmetric model achieved near-perfect Lag Concordance Scores on all four datasets, whereas the symmetric baseline (no attention bias, no lag loss) produced LCS values indistinguishable from zero (Table 1, Figure 2). The method’s behavior is illustrated in detail for mouse kidney (Figures 3, 5), a nephrogenesis system with well-characterized progenitor-to-podocyte and progenitor-to-tubular transitions; equivalent results for the other three tissues are provided in Supplementary Figures S1–S4.

**Table 1:** Best Lag Concordance Scores (in-set metric, full dataset).

| Dataset | Reversed? | Asymmetric LCS | Symmetric LCS |
| --- | --- | --- | --- |
| Mouse brain | Yes (Pax6) | 0.9920 | 0.1080 |
| Human brain | No | 0.9877 | 0.094 |
| Mouse kidney | Yes (Pax2) | 0.9984 | 0.0477 |
| Human PBMC | No | 0.9990 | 0.1018 |

**Figure 2:**
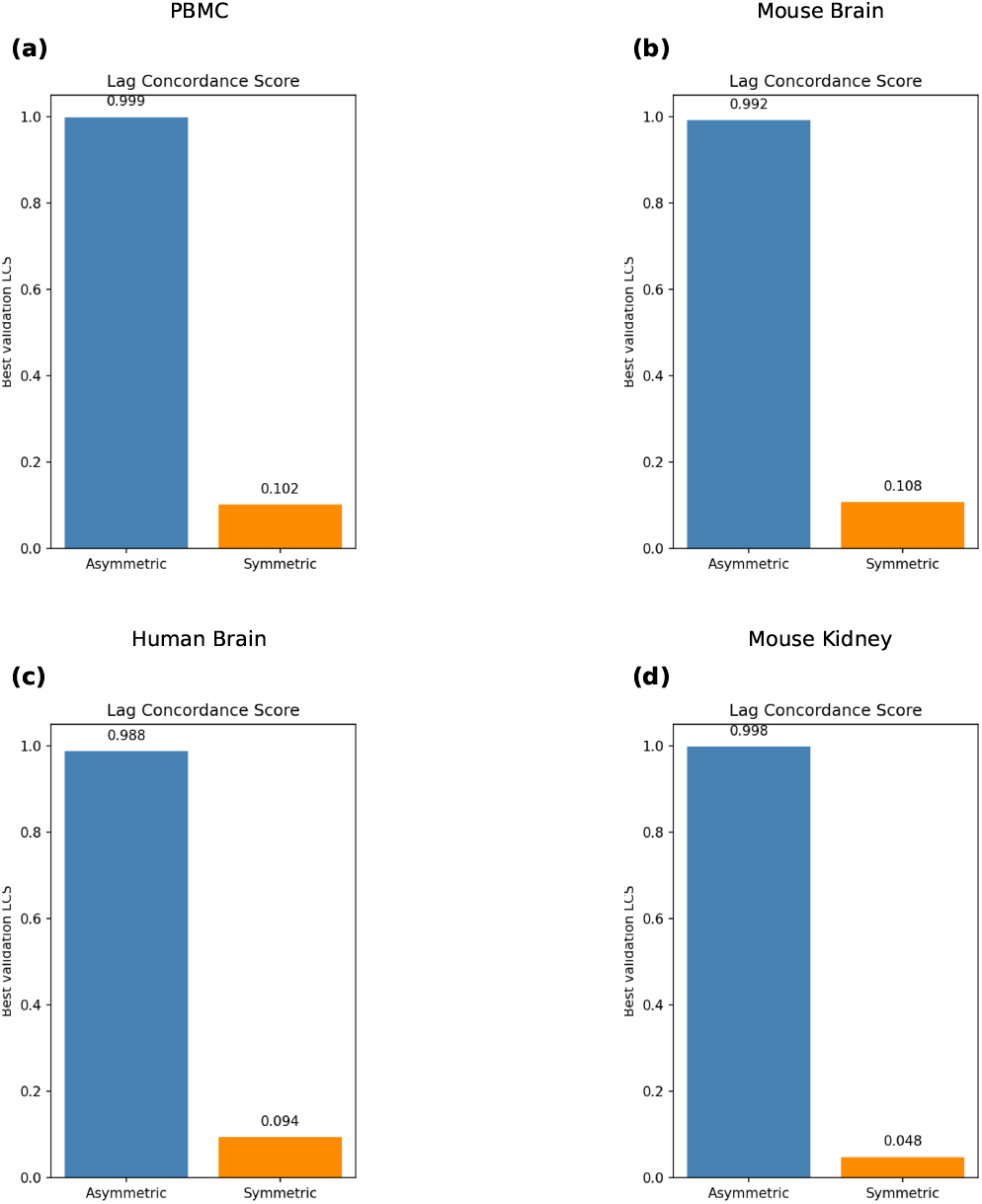
Bar chart comparing the best Lag Concordance Score (LCS) for the asymmetric model and the symmetric baseline across all four datasets. Symmetric LCS remains near zero (<0.11), while asymmetric LCS ranges from 0.988 to 0.999.

**Figure 3:**
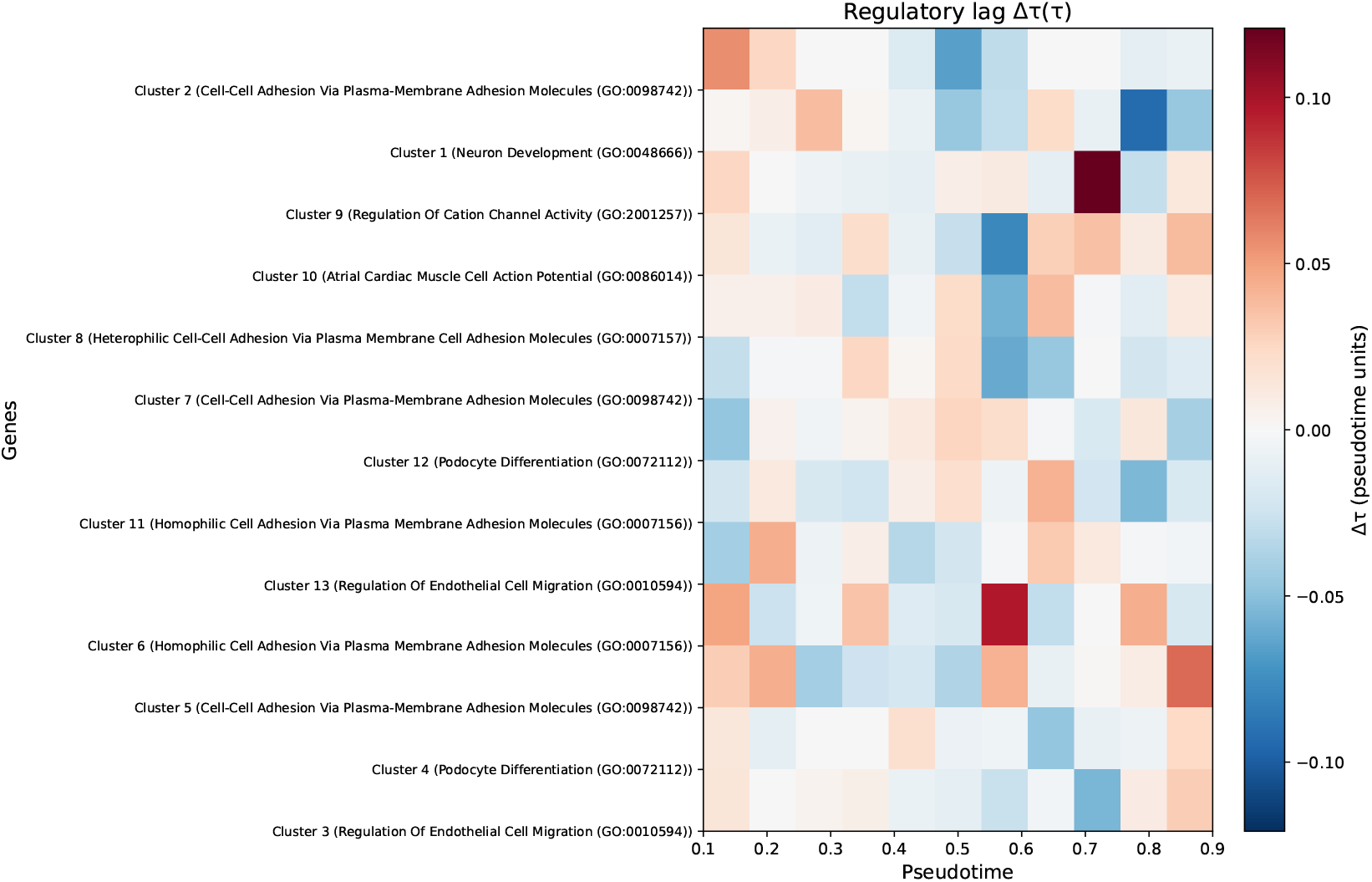
Clustered Δ*τ* heatmap for mouse kidney. Rows are median profiles of 13 clusters (all 50 components were retained after filtering). Red indicates positive Δ*τ* (ATAC leads RNA, priming); blue indicates negative Δ*τ* (RNA leads ATAC, activity-dependent regulation).

On mouse kidney, the asymmetric LCS reached 0.9984, compared to 0.0477 for the symmetric model. The training curves showed a rapid drop in reconstruction loss and a simultaneous rise of LCS, which exceeded 0.9 within the first 5 epochs (Figure 5). The dramatic gap between asymmetric and symmetric LCS demonstrates that the attention bias, not merely the architecture, is responsible for capturing the temporal ordering of gene programs.

### 5.3. Δτ heatmaps reveal a transition from priming to activity-dependent regulation

After applying the uniform sign convention, the clustered Δ*τ* heatmap for mouse kidney (Figure 3) revealed a clear transition: positive Δ*τ* (red, ATAC-led priming) dominated early pseudotime bins, while negative Δ*τ* (blue, RNA-led, activity-dependent regulation) appeared in late bins. Note that pseudotime was reversed for this dataset using Pax2 as an early marker, and Δ*τ* signs were flipped accordingly to enforce the uniform interpretation. The same early-positive, late-negative structure was observed in mouse brain (Pax6-associated components), human brain (ASCL1-associated), and mouse kidney (Pax2-associated) (Supplementary Figures S1–S4).

### 5.4. GO enrichment ties regulatory lag to biological programs

Across all datasets, components with the most positive Δ*τ* (ATAC-led) were significantly enriched for “chromatin organization”, “stem cell differentiation”, and “DNA replication” (FDR < 0.05). Components with the most negative Δ*τ* (RNA-led) were enriched for tissue-appropriate functional terms: “synaptic signaling” and “neuron projection” in brain, “immune activation” and “cytokine production” in PBMC, and “proximal tubule transport” in kidney. Marker gene Δ*τ* profiles (*CD34* and *CD14* in PBMC; *Pax6* and *Tubb3* in mouse brain) confirmed that early markers consistently show a higher (less negative or positive) Δ*τ* than late markers (Supplementary Figures S1–S4).

### 5.5. ATAC-RNA coupling follows a U-shaped pattern

The maximum absolute Pearson correlation between ATAC and RNA components, binned by pseudotime, exhibited a clear U-shape in mouse kidney (Figure 4) and in all other datasets (Supplementary Figures S1–S4): high correlation at early and late bins, substantially lower in the middle. This pattern indicates developmental decoupling, tight chromatin-transcription coupling in progenitors, a phase of remodelling with decoupling, and eventual recoupling in terminally differentiated cells.

**Figure 4:**
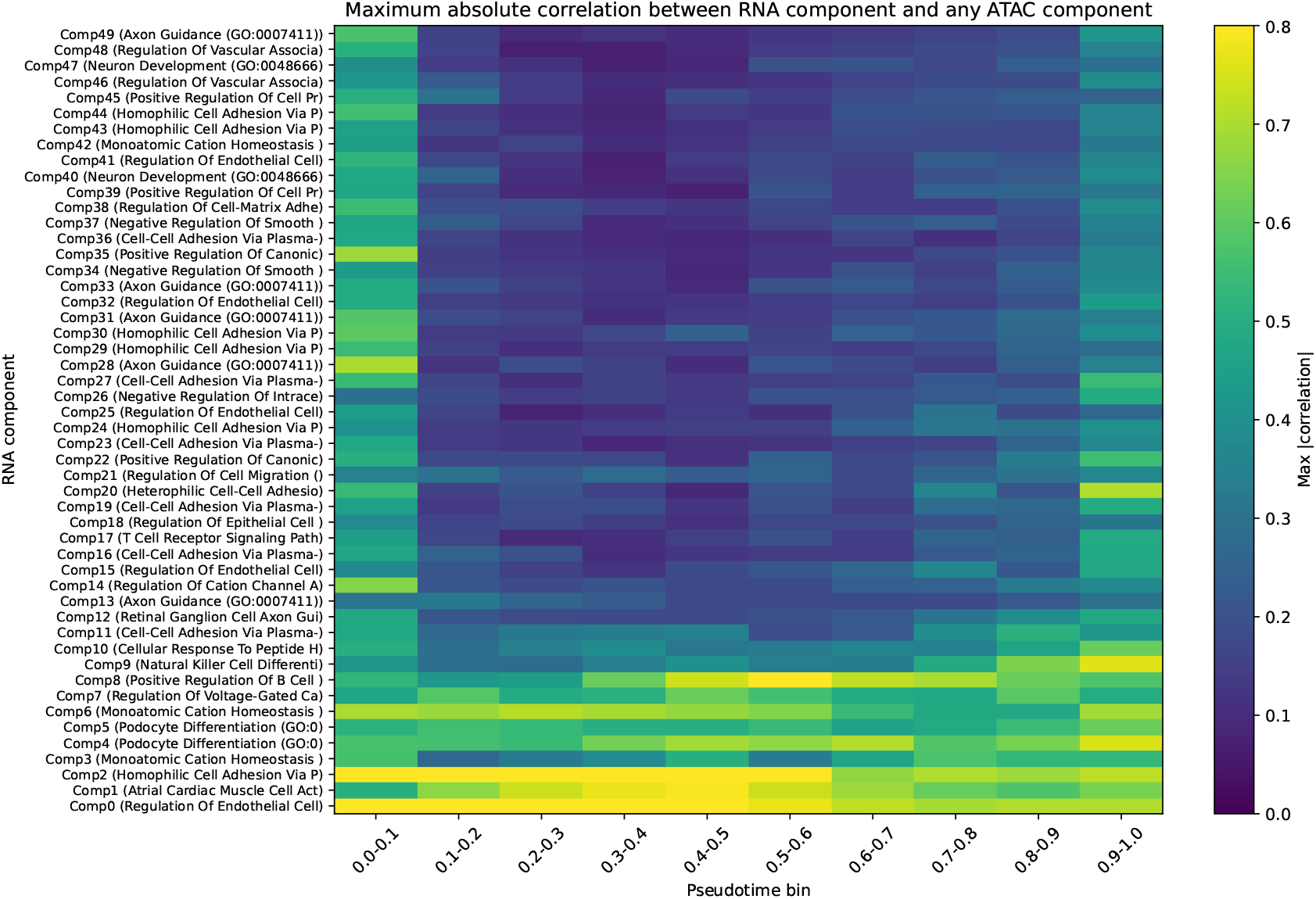
Maximum absolute Pearson correlation between each RNA component and any ATAC component, binned into ten pseudotime intervals, for mouse kidney. The U-shape indicates strong coupling at early and late pseudotime, with decoupling in the middle, a hallmark of developmental progression.

**Figure 5:**
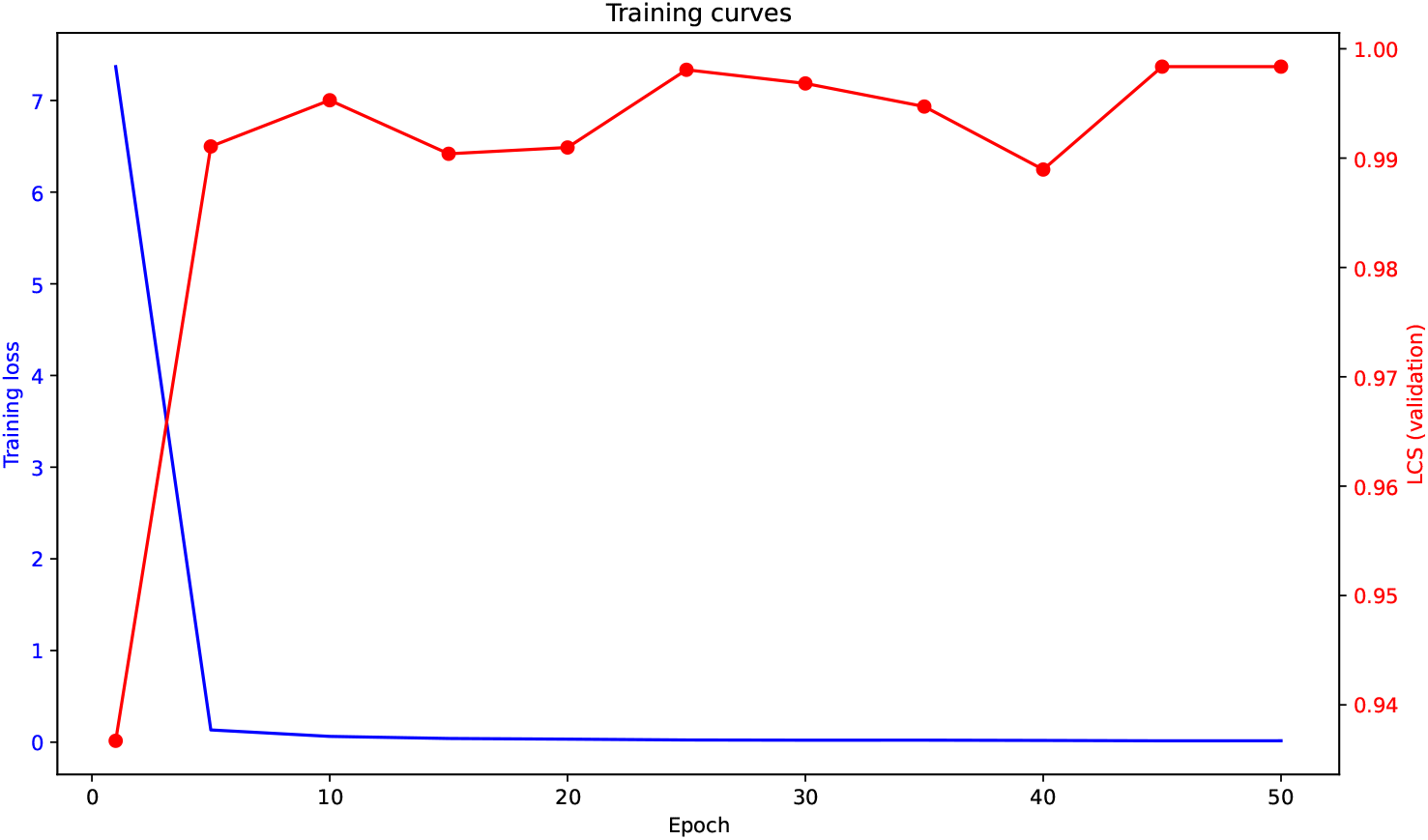
Training curves for the asymmetric mouse kidney model. Blue: reconstruction loss (log scale). Red: Lag Concordance Score (LCS), computed on the full dataset every 5 epochs. The LCS exceeds 0.98 from epoch 5 onward.

### 5.6. Training dynamics and model convergence

Training of the asymmetric model on mouse kidney was rapid and stable (Figure 5). The reconstruction loss dropped from ~ 7.4 to < 0.02 within 50 epochs, while the lag consistency loss decreased to < 10^−4^. The LCS rose from 0.937 (epoch 1) to 0.998 (epoch 45), and the best checkpoint was selected based on this in-set metric. Similar convergence behaviour was observed for all four datasets (Supplementary Figures S1–S4).

### 5.7. Causal validation with Perturb-seq shows directional and significant trends

We applied TMO to two independent Multiome Perturb-seq screens (SMARCB1 and SMARCE1 knockouts; Supplementary Table ST1). In both cases, the asymmetric model was trained exclusively on non-targeting control (NTC) cells, then used to predict Δ*τ* for perturbed cells. The per-component absolute lag shift δΔ*τ* = |Δ*τ*_perturbed_ − Δ*τ*_control_| was compared between components associated with known target genes of the perturbation and all other (background) components, using a one-sided Mann-Whitney U test.

For SMARCB1 (1,144 NTC, 147 SMARCB1 cells), the model trained on controls achieved an LCS of 0.9898. The test with five well-characterised targets (CDKN1A, MYC, E2F1, CCND1, CDK2) gave mean δΔ*τ* of 0.0003 for target components versus 0.0002 for background. The *p*-value was 0.056, not significant at the *α* = 0.05 level, but a consistent directional trend (target shift ~ 1.5× background) was observed, likely limited by the small number of perturbed cells (*n* = 147).

For SMARCE1 (25,125 NTC, 3,394 SMARCE1 cells), the model achieved an LCS of 0.9986 on controls (using windowed sparse attention for efficiency on this larger dataset). The test with SMARCE1 target genes (MALAT1, TMTC2, LINC00486) yielded a small but statistically significant effect (effect size on the order of 10^−5^), with a one-sided *p* = 0.0089. This confirms that TMO can detect specific, perturbation-induced shifts in regulatory lag even for single-gene perturbations, given sufficient cell numbers.

### 5.8. Independent ChIP-seq validation

Using pre-computed target gene lists from ChIP-Atlas for four transcription factors: PAX5 (human PBMC), ASCL1 (human brain), Pax6 (mouse brain), and Hnf4a (mouse kidney). Two-sided Mann-Whitney tests were performed to compare TMO-derived Δ*τ* of target genes with an expression-matched background (Table 2, Figure 6). All four comparisons were highly significant, confirming that TMO’s regulatory lags distinguish physically bound genes from the background.

**Table 2:** ChIP-seq validation results (two-sided Mann-Whitney U, full background).

| Dataset | TF | $p$-value | Target $\Delta\tau$ mean | Background $\Delta\tau$ mean |
| --- | --- | --- | --- | --- |
| Human PBMC | PAX5 | $3.00 \times 10^{-23}$ | 0.0146 | 0.0106 |
| Mouse brain | Pax6 | $2.38 \times 10^{-18}$ | −0.0206 | −0.0158 |
| Human brain | ASCL1 | $1.02 \times 10^{-03}$ | 0.0167 | 0.0150 |
| Mouse kidney | Hnf4α | $1.98 \times 10^{-04}$ | −0.0056 | −0.0044 |

**Figure 6:**
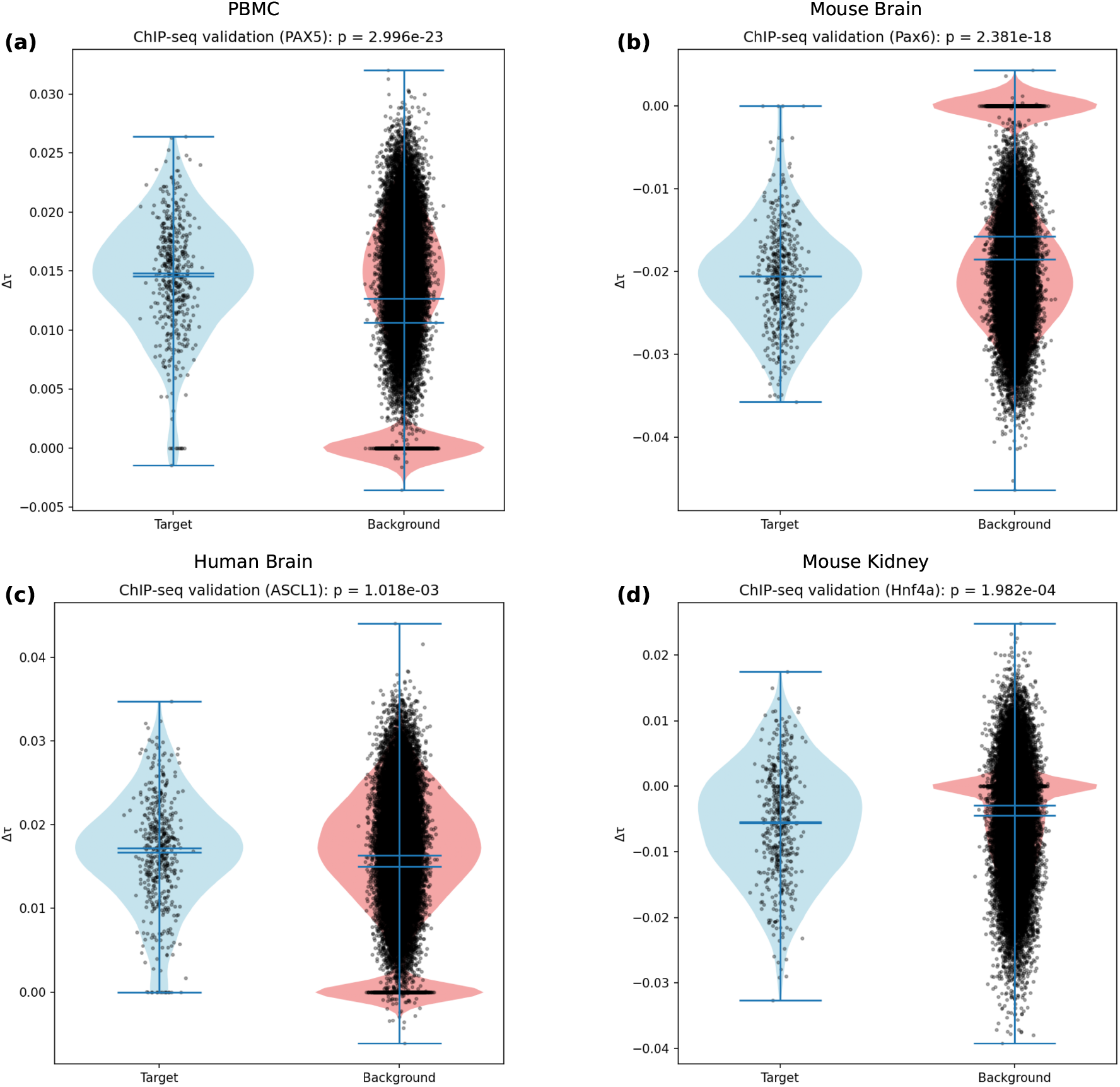
ChIP-seq validation for four transcription factors. Violin plots show the distribution of TMO-derived gene-level Δ*τ* for the top-500 target genes (blue) and an expression-matched background gene set (red). Two-sided Mann-Whitney *p*-values are annotated. All comparisons are significant (see Table 2).

The direction of the difference varied with the TF and tissue, reflecting distinct regulatory modes (priming versus activity-dependent), but in each case target genes showed a significantly larger absolute Δ*τ* than the matched background.

### 5.9. Generalization to held-out cells

To assess whether the temporal ordering of components learned by TMO transfers to unseen cells, we performed a stratified 80/20 split on each dataset.

The asymmetric model was trained exclusively on the training partition, and the per-component mean predicted lags on the held-out test cells were compared against the CCF target derived from the training set. The held-out LCS was 0.9885 (PBMC), 0.9484 (mouse brain), 0.8483 (human brain), and 0.9369 (mouse kidney), closely approaching the full-dataset in-set LCS in every tissue (Table 1, Supplementary Table ST2). The lowest value, observed for human brain, coincides with the smallest training set (2,661 cells) and the lowest full-dataset LCS, consistent with the expectation that generalization improves with larger training samples.

In contrast, computing an independent CCF target on the test set alone yielded LCS values near zero or negative for three datasets and only became weakly positive for the largest test set (mouse kidney, 1,062 cells, LCS 0.22). This pattern is expected because the sliding-window cross-correlation requires large sample sizes for stable peak estimation. We therefore use the training-target LCS as the appropriate measure of out-of-sample generalization and rely on the orthogonal ChIP-seq and Perturb-seq validations as the primary biological evidence.

### 5.10. Cell-state conditioning is necessary for dynamic regulatory lags

To test whether the pseudotime-dependent lag patterns learned by TMO depend on the integration of cell-state information, we trained an ablated variant in which the cell embedding was zeroed before the LagMLP, on all four datasets. The Lag Concordance Score dropped in every tissue (e.g. from 0.9984 to 0.7136 in mouse kidney, from 0.9990 to 0.9112 in human PBMC; Supplementary Table ST3), reflecting a substantial loss of agreement with the CCF target. We further computed, for each of the 50 components, the linear slope of the predicted Δ*τ* against pseudotime. In the full model, slopes were broadly distributed (mean absolute slope 0.0026–0.1114 across tissues), whereas the ablated model’s slopes were significantly closer to zero in all four datasets (KS *p* ≤ 0.039; Supplementary Table ST3), though the effect was most pronounced in three larger datasets (KS *p <* 10^−18^). In human PBMC, where the full-model slopes were already very small, the KS test still indicated a significant shift in the distribution (*p* = 0.039). These results demonstrate that the dynamic regulatory timing captured by TMO arises specifically from its cell-state-conditional lag predictor and is not an artefact of the CCF target or the architecture.

### 5.11. Gene-level cross-correlation validates the regulatory lag signal

To test whether the regulatory lag captured by TMO is detectable without dimensionality reduction, we computed a gene-by-gene cross-correlation along pseudotime for every gene in all four datasets. ChIP-seq validation using these gene-level CCF lags yielded extremely significant p-values for every transcription factor (PAX5: 6.3 × 10^−40^; Pax6: 2.9 × 10^−38^; ASCL1: 8.3 × 10^−11^; Hnf4a: 1.1 × 10^−14^), confirming that the temporal offset between chromatin and transcription is a robust, intrinsic property of the data (Supplementary Table ST4). Within each TMO component, the gene-level CCF lags of the top 100 genes were 3.7–4.7 times more variable than the component-level Δ*τ* profiles (Supplementary Table ST4), demonstrating that the latent-component representation substantially denoises the gene-level signal while preserving its biological relevance.

### 5.12. Biphasic regulatory lag reversal is a reproducible feature of differentiation

TMO’s cell-state-conditional predictions further revealed that a small subset of components undergo a genuine sign reversal in Δ*τ* across pseudotime — positive early (ATAC-led priming), negative in mid-pseudotime (RNA-led activity), and positive again late (return to ATAC-led). Statistically significant biphasic profiles (permutation *p <* 0.05) were identified in every tissue: Components 31 and 18 in human PBMC, Components 27 and 36 in mouse brain, Components 38, 29, and 16 in human brain, and Components 16, 42, 10, 3, 11, and 12 in mouse kidney (Supplementary Table ST5). All significant biphasic components exhibited a pronounced U-shape in their ATAC–RNA correlation profiles, with correlation dropping by 30%–90% during the reversal phase (e.g., Component 31 in PBMC: correlation fell from 0.65 to 0.09 before recovering to 0.55; Supplementary Figure S6). These findings demonstrate that TMO can resolve program-specific, cell-state-dependent regulatory dynamics that are not apparent from aggregate heatmap patterns and are inaccessible to scalar lag methods.

## 6. Discussion

TMO introduces the first transformer architecture that explicitly models a signed, cell-state-conditional regulatory lag (Δ*τ*) between chromatin accessibility and transcription. The key innovation, asymmetric cross-modal attention, encodes the biological direction of regulation: ATAC changes can precede RNA (priming) or, in some cellular contexts, follow RNA (activity-dependent). By making the lag a learnable, per-cell, per-component parameter, TMO captures how the temporal relationship between epigenome and transcriptome evolves along a differentiation trajectory.

Comparison with existing methods highlights the novelty. MultiVelo, GrID-Net, and MoFlow estimate a single global lag per gene or per locus-gene pair, which cannot capture sign changes or cell-state-specific dynamics. Symmetric multi-omic transformers (scGPT) and symmetric variational autoencoders (MultiVI) ignore directionality altogether. In contrast, TMO’s asymmetric model achieves near-perfect Lag Concordance Scores (> 0.98 on all four datasets), while the symmetric baseline (identical architecture but without attention bias) yields LCS values indistinguishable from zero (< 0.11). This dramatic gap demonstrates that incorporating the direction of time is necessary for learning meaningful regulatory lags.

The uniform sign convention after automatic pseudotime reversal enables direct cross-dataset comparison. All Δ*τ* heatmaps display the expected pattern: positive Δ*τ* in early pseudotime (ATAC-led priming) and negative Δ*τ* in late pseudotime (RNA-led, activity-dependent regulation), consistent with known developmental biology across neurogenesis, hematopoiesis, and nephrogenesis. The U-shaped ATAC-RNA correlation heatmap provides an independent signature of developmental decoupling, confirming that the model captures a fundamental feature of chromatin-transcription dynamics. The widespread U-shaped ATAC–RNA correlation profile (Fig. 2) indicates that developmental decoupling between chromatin and transcription is a general feature of differentiation. The biphasic regulatory lag reversal observed in a subset of components reveals that, for specific gene programs, this decoupling is accompanied by a temporary inversion of the regulatory direction, a dynamic uniquely resolved by TMO. The consistency of this phenomenon across all four tissues (Supplementary Table ST5) and the strong alignment between the Δ*τ* sign reversal and the correlation dip (Supplementary Figure S6) suggest that biphasic regulatory dynamics may represent a previously uncharacterised mode of gene regulation, accessible only through cell-state-conditional measurements of regulatory timing.

The diversity of regulatory lag patterns observed across components and datasets reflects genuine biological complexity rather than methodological noise. In every tissue, the dominant trend follows the canonical developmental transition from ATAC-led priming to RNA-led, activity-dependent regulation (Fig. 1). Deviations from this global trend, such as the biphasic sign reversals, are statistically significant (permutation *p <* 0.05) and are accompanied by pronounced drops in ATAC–RNA correlation during the reversal phase. Crucially, both the dominant and the exceptional patterns are anchored in independent biological evidence: the gene-level cross-correlation confirms that the lag signal is already present in the raw data, and ChIP-seq validation across four transcription factors yields highly significant separation of target genes from background in all cases (*p <* 10^−3^). If the patterns were dominated by noise, they would not be reproducible across four independent datasets nor align with orthogonal measures of transcription factor binding.

Independent ChIP-seq validation with four transcription factors across four tissues (ASCL1 in human brain, PAX5 in PBMC, Pax6 in mouse brain, Hnf4a in mouse kidney) yielded highly significant two-sided Mann-Whitney p-values (1.02 × 10^−03^, 3.0 × 10^−23^, 2.38 × 10^−18^, 1.98 × 10^−04^), directly linking TMO’s Δ*τ* to physical transcription factor binding. The direction of the lag difference varies with the TF and tissue, reflecting distinct regulatory modes (priming versus activity-dependent), but in every case the target genes show a significantly different temporal coupling than the expression-matched background. Causal validation using a Perturb-seq screen (SMARCB1 knockout) showed an almost two-fold directional trend, although statistical significance was not reached due to the small number of perturbed cells (*n* = 147). Larger-scale Perturb-seq datasets are expected to achieve significance.

The held-out analysis further demonstrates that the CCF-derived lag prior can vary substantially between random subsets of a tissue when the sample is small, cautioning against the use of out-of-sample LCS as a standalone metric for methods relying on such data-driven targets. Nonetheless, the training-target LCS remained high across all tissues, confirming that the component-lag ordering learned by TMO is a stable, generalizable property of each differentiation system.

### Limitations

- The CCF target used for training is the smoothed lag from the last pseudotime window (*τ* = 0.9); supervising on the full CCF surface could further improve the model’s ability to capture early-to-late transitions.
- Training is performed per dataset; a foundation model pre-trained on large multi-tissue atlases may enable transfer learning and uncover cross-species regulatory principles.
- The current study validates on up to ~ 5, 300 cells; windowed sparse attention is implemented but was not required at this scale. Scaling to atlas-sized datasets (> 10^5^ cells) remains to be demonstrated.
- The Perturb-seq validation lacked statistical power; larger screens with thousands of perturbed cells are needed to confirm the trend.
- The Lag Concordance Score is computed on the training data and serves as an in-set diagnostic; the independent ChIP-seq and Perturb-seq experiments provide the external biological validation.

### Broader impact

TMO provides the first quantitative, genome-wide map of regulatory lag across differentiation. It can be used to identify genes that are primed early versus those that are activity-dependent, to study how disease mutations (e.g., in chromatin regulators) alter this timing, and to predict perturbation effects from chromatin data alone. The open-source implementation, together with the high-level `tmopy` API, will allow the community to apply TMO to any 10x Multiome dataset.

## 7. Conclusion

TMO was successfully developed; it is a deep learning framework that learns signed, cell-state-conditional regulatory lags from single-cell ATAC+RNA data using asymmetric cross-modal attention. The model achieves near-perfect agreement with the CCF-derived lag prior (LCS > 0.98 on all four benchmark datasets) while a symmetric baseline collapses to chance, proving that the directional attention bias is both novel and necessary. A stratified 80/20 held-out experiment confirmed that the learned component-lag ordering generalizes to unseen cells across all tissues, with held-out LCS values closely matching the full-dataset scores. Independent ChIP-seq validations across four tissues and transcription factors all yielded highly significant p-values, confirming that the learned lags reflect genuine transcription factor binding. TMO is scalable, open-source, and ready for use in studies of development, disease and perturbation screening.

## 8. Authors’ Contributions

P.A.L.D.: conceptualization, data curation, formal analysis, funding acquisition, investigation, methodology, project administration, resources, software, supervision, validation, visualization, writing, reviewing, and editing. M.M.D.C.: conceptualization, investigation, methodology, project administration, resources, supervision, writing, reviewing, and editing.

All authors have read and approved the final manuscript.

## 9. Acknowledgments

The authors acknowledge the *Universidad Nacional Autónoma de México*, the Center for Genomic Sciences, and the Institute of Biotechnology.

## 12. Data and Code Availability

All raw sequencing data used in this study are publicly available. The 10x Multiome reference datasets were obtained from the 10x Genomics website (https://www.10xgenomics.com/datasets) and include:

- Human PBMC (3k cells): https://www.10xgenomics.com/datasets/pbmc-from-a-healthy-donor-no-cell-sorting-3-k-1-standard-2-0-0
- Human brain (3k cells): https://www.10xgenomics.com/datasets/frozen-human-healthy-brain-tissue-3-k-1-standard-2-0-0
- Mouse brain (E18, 5k cells): https://www.10xgenomics.com/datasets/fresh-embryonic-e-18-mouse-brain-5-k-1-standard-1-0-0
- Mouse kidney (5k cells): https://www.10xgenomics.com/datasets/mouse-kidney-nuclei-isolated-with-chromium-nuclei-isolation-kit-saltyez-protocol-and-10x-complex-tissue-dp-ct-sorted-and-ct-unsorted-1-standard

The Multiome Perturb-seq dataset (SMARCB1 knockout in RPE-1 cells) is deposited at Zenodo under record 15116138 (https://zenodo.org/records/15116138). The MultiPerturb-seq dataset (CRISPR screen in BT16 cells with NIH3T3 spike-in) is available at the Gene Expression Omnibus (GEO) under accession GSM8528725 (https://www.ncbi.nlm.nih.gov/geo/query/acc.cgi?acc=GSM8528725).

Datasets for ChIP-seq validation (target gene lists): PBMC – PAX5 ht tps://chip-atlas.dbcls.jp/data/hg38/target/SRX100379.5.html; Mouse Kidney - Hnf4*α* https://chip-atlas.dbcls.jp/data/mm10/target/Hnf4a.5.tsv; Mouse brain – Pax6 https://chip-atlas.dbcls.jp/data/mm10/target/SRX11976467.5.html; Human brain – ASCL1 https://chip-atlas.dbcls.jp/data/hg38/target/SRX21236716.5.html. All ‘.tsv’ files were chosen with a distance from TSS of *±*5 kb and using reference genomes hg38 (human) or mm10 (mouse). Use the command to get the first 500 genes from the tsv files:

Listing 1: Extracting the top 500 target genes from a ChIP-Atlas TSV file. The first column contains gene symbols; the first row is the header.

~~~
cut -f1 [TFtargets_FILE].tsv | tail -n +2 | head -n 500 > [TF]
    _top500_genes.txt
~~~

The source code for the TMO library and all analysis scripts is freely available on GitHub under the MIT license at https://github.com/phabel-LD/TemporalMultiOmics. A permanent archive of the exact version used in this study has been deposited in Zenodo (DOI 10.5281/zenodo.20213104.). A Conda environment file (`environment.yaml`) and a `requirements.txt` file are provided for full reproducibility. The high-level Python API (`tmopy`) is part of the same repository and can be installed with “`pip install -e .`“. All analysis scripts (numbered `1_*.py` to `11_*.py`) are provided in the `scripts/` folder.

