## Supplementary Material for "TMO: ASYMMETRIC CROSS-MODAL ATTENTION FOR LEARNING CELL-STATE-DEPENDENT REGULATORY LAGS FROM SINGLE-CELL MULTIOMIC DATA"

#### S1 – Mouse brain

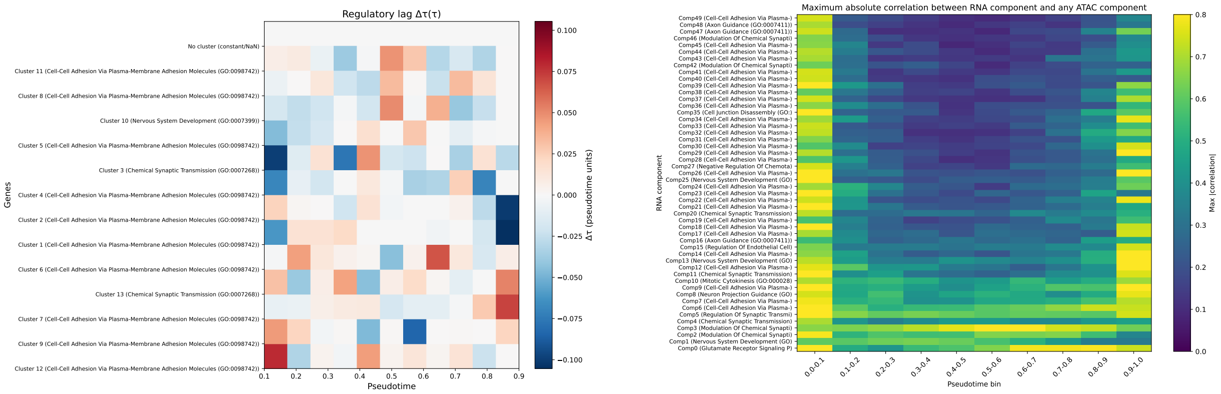


Figure 1: Mouse brain: $\Delta\tau$ heatmap (left) and ATAC-RNA correlation U-shape (right).

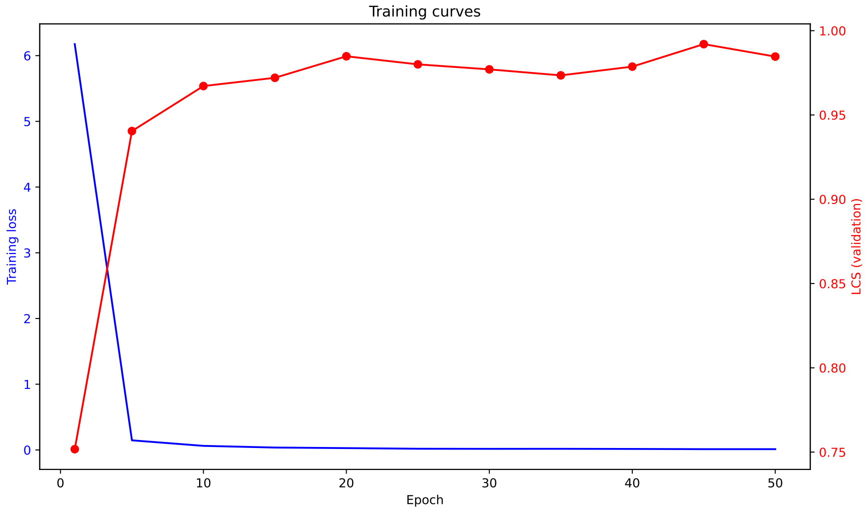


Figure 2: Mouse brain: training curves (reconstruction loss + LCS).

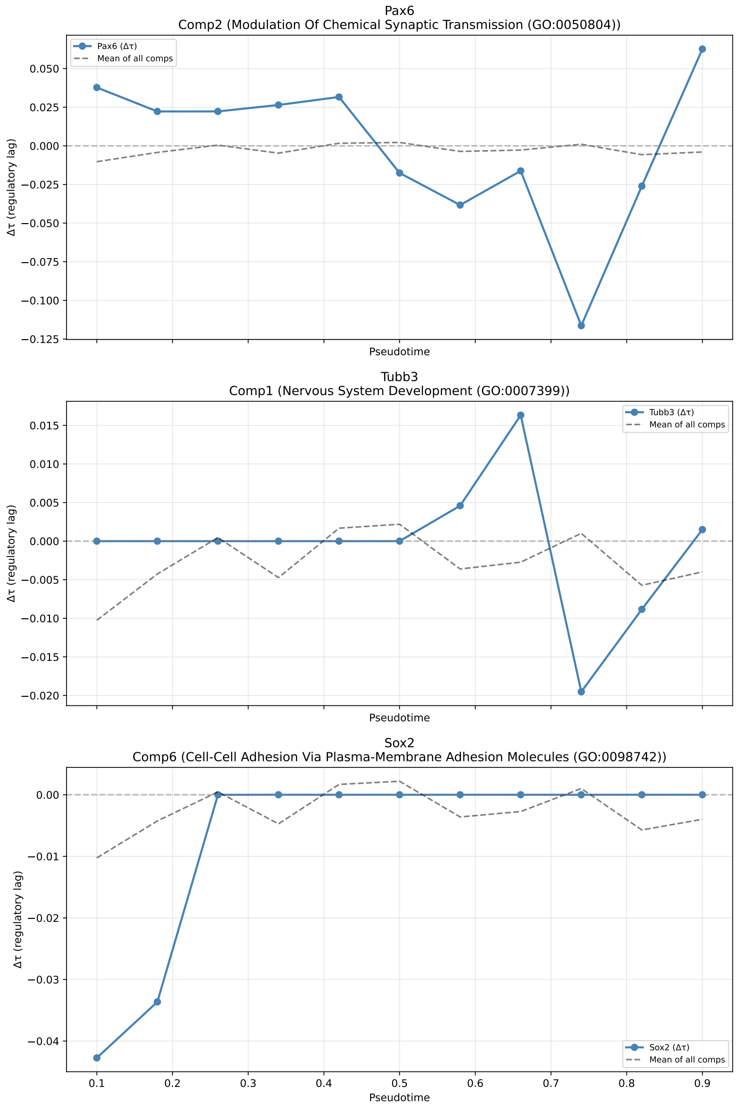


Figure 3: Mouse brain: marker $\Delta\tau$ profiles (Pax6, Tubb3, Sox2).

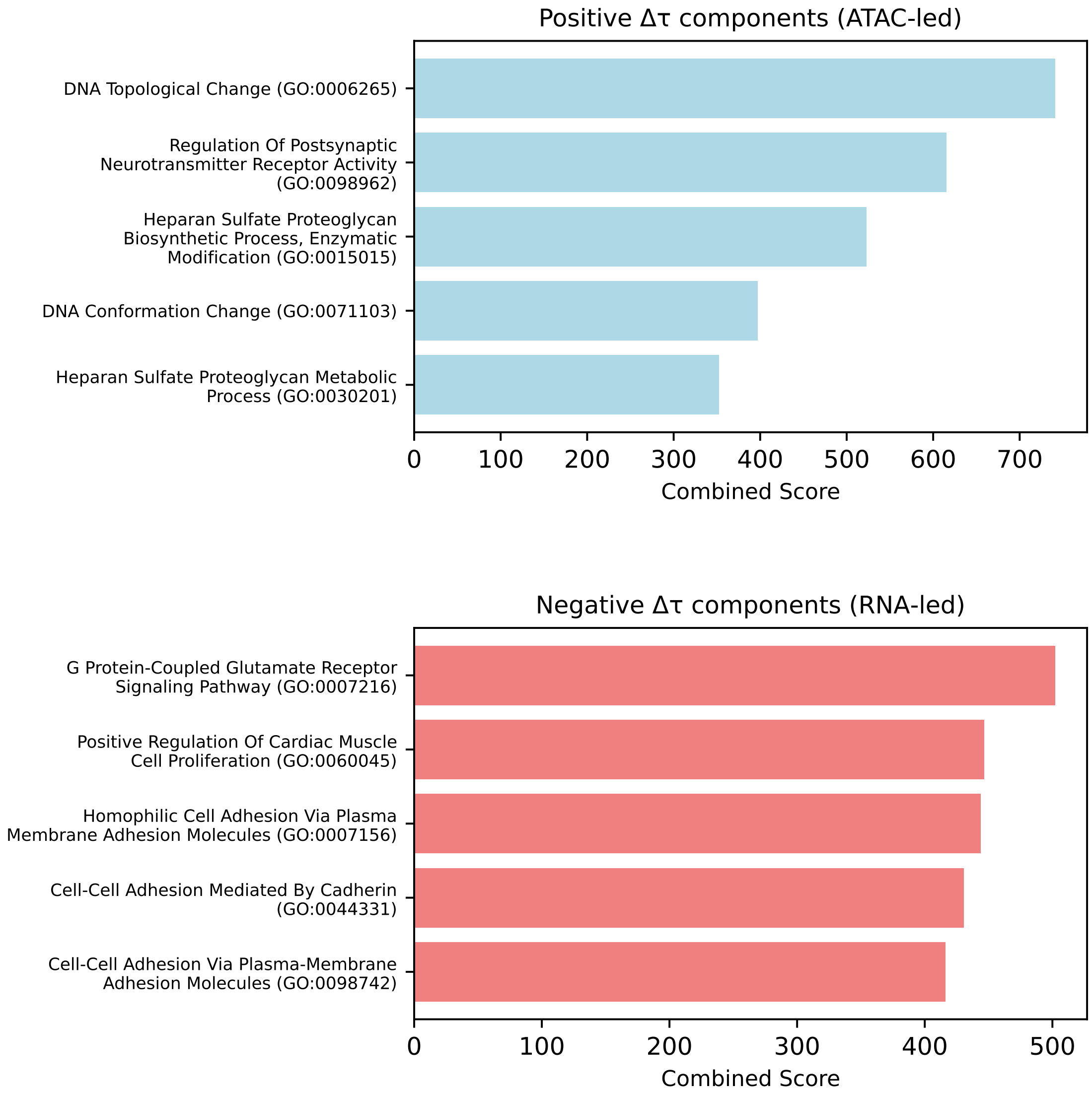


Figure 4: Mouse brain: Gene-program GO enrichment.

### S2 – Human brain

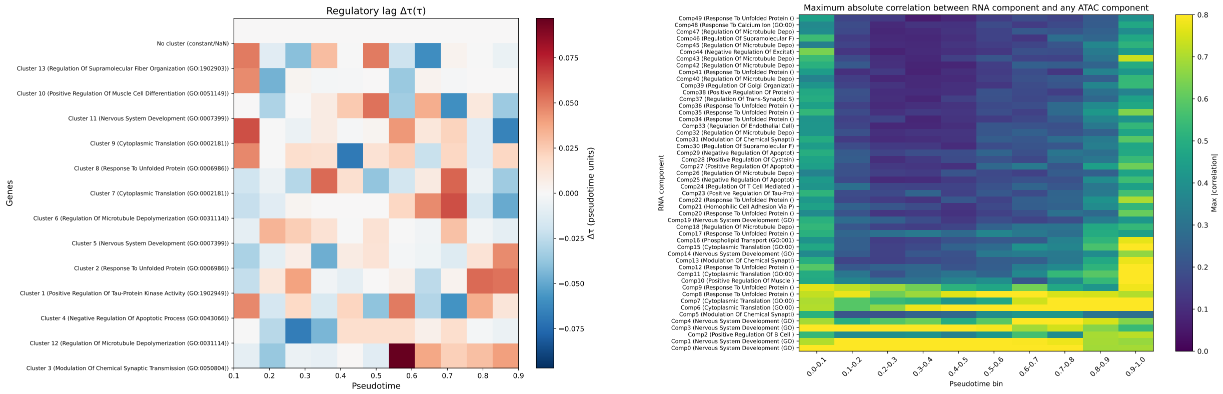


Figure 5: Human brain: $\Delta\tau$ heatmap (left) and ATAC-RNA correlation U-shape (right).

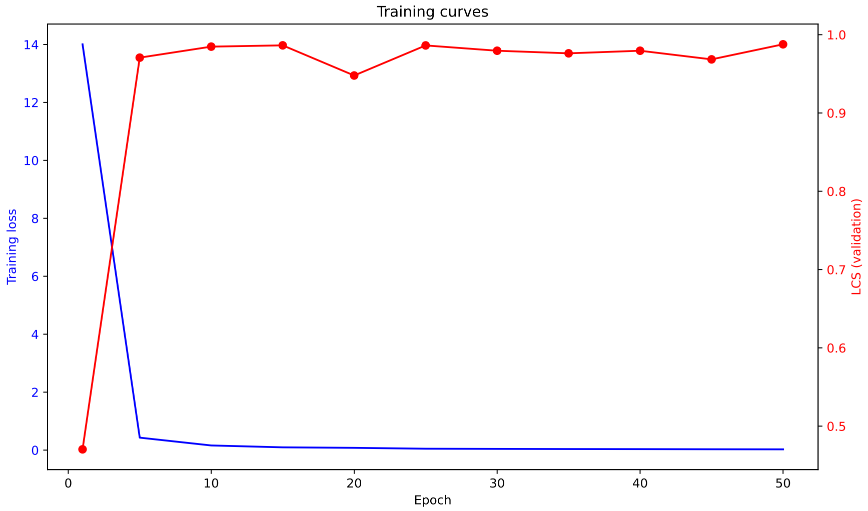


Figure 6: Human brain: training curves (reconstruction loss + LCS).

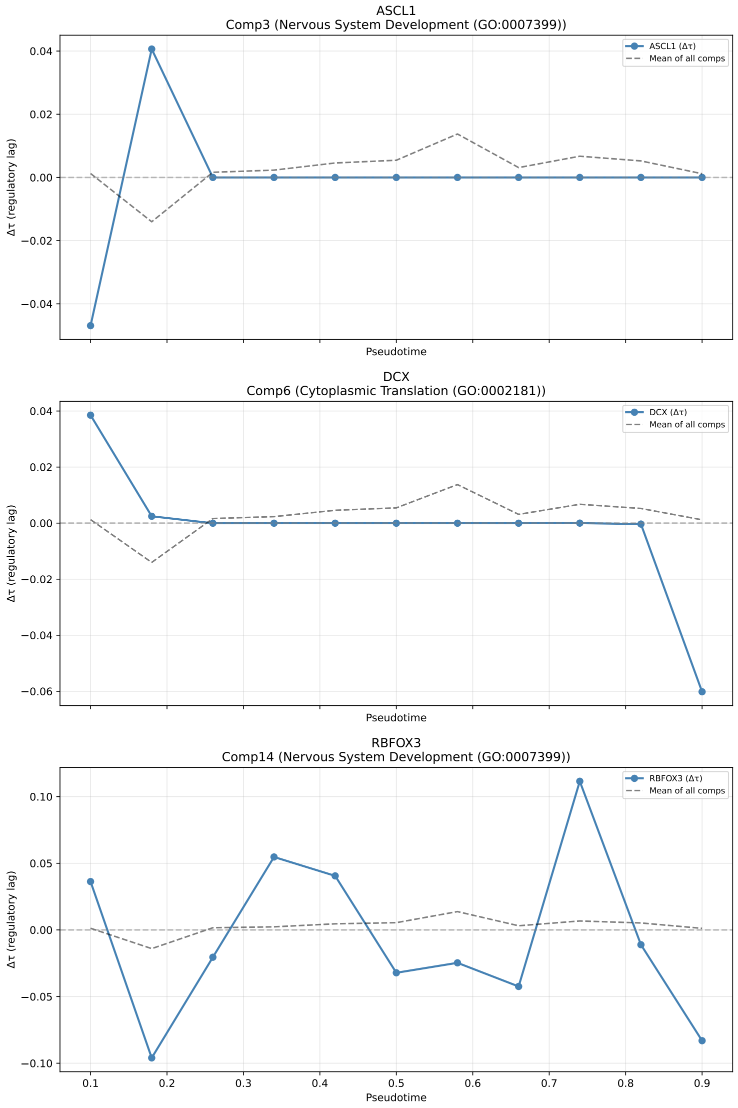


Figure 7: Human brain: marker $\Delta\tau$ profiles (ASCL1, DCX, RBFOX3).

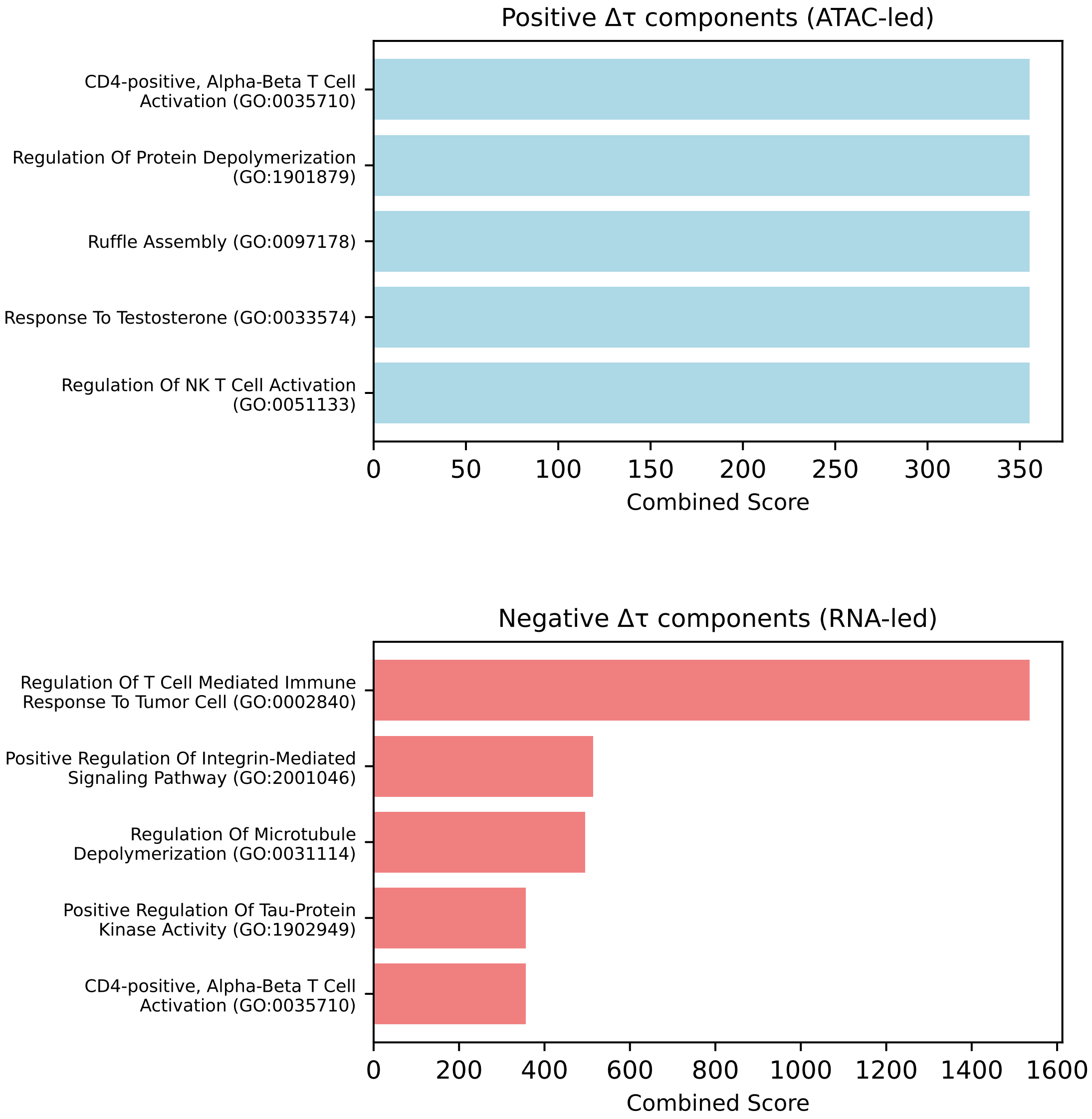


Figure 8: Human brain: Gene-program GO enrichment.

#### S3 – Human PBMC

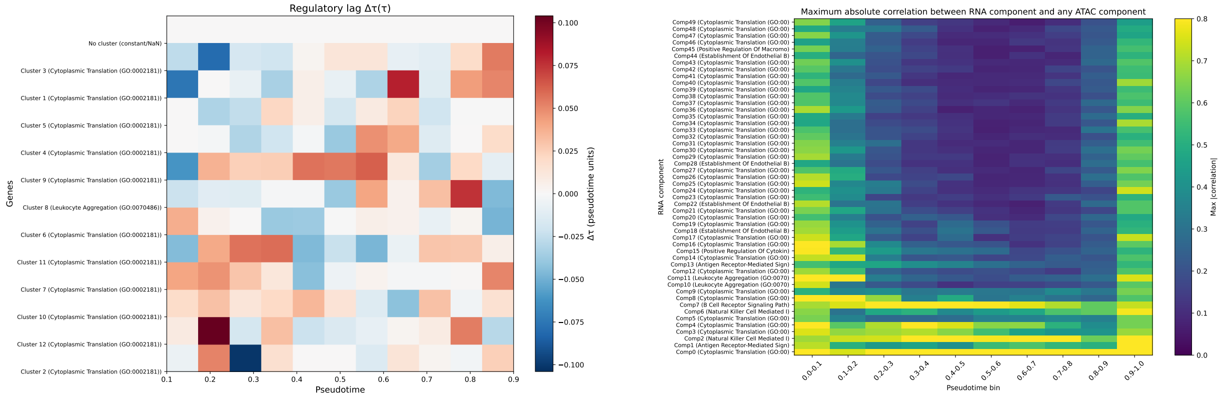


Figure 9: Human PBMC: $\Delta\tau$ heatmap (left) and ATAC-RNA correlation U-shape (right).

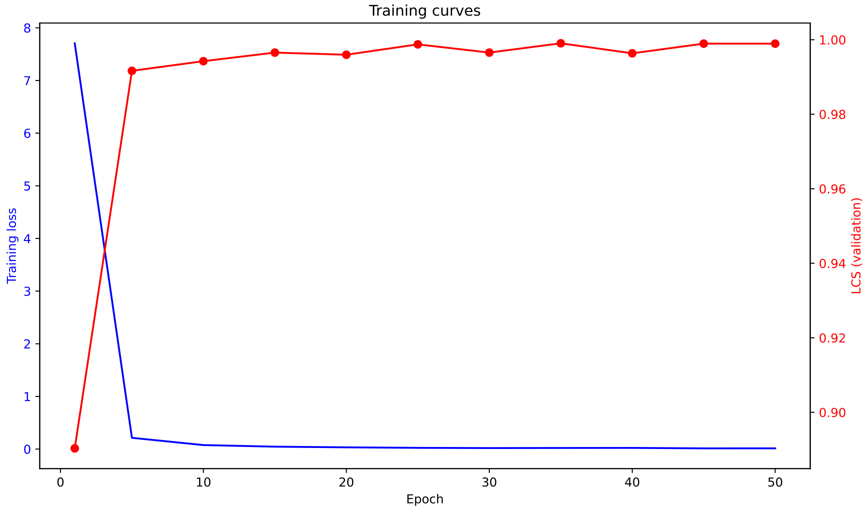


Figure 10: Human PBMC: training curves (reconstruction loss + LCS).

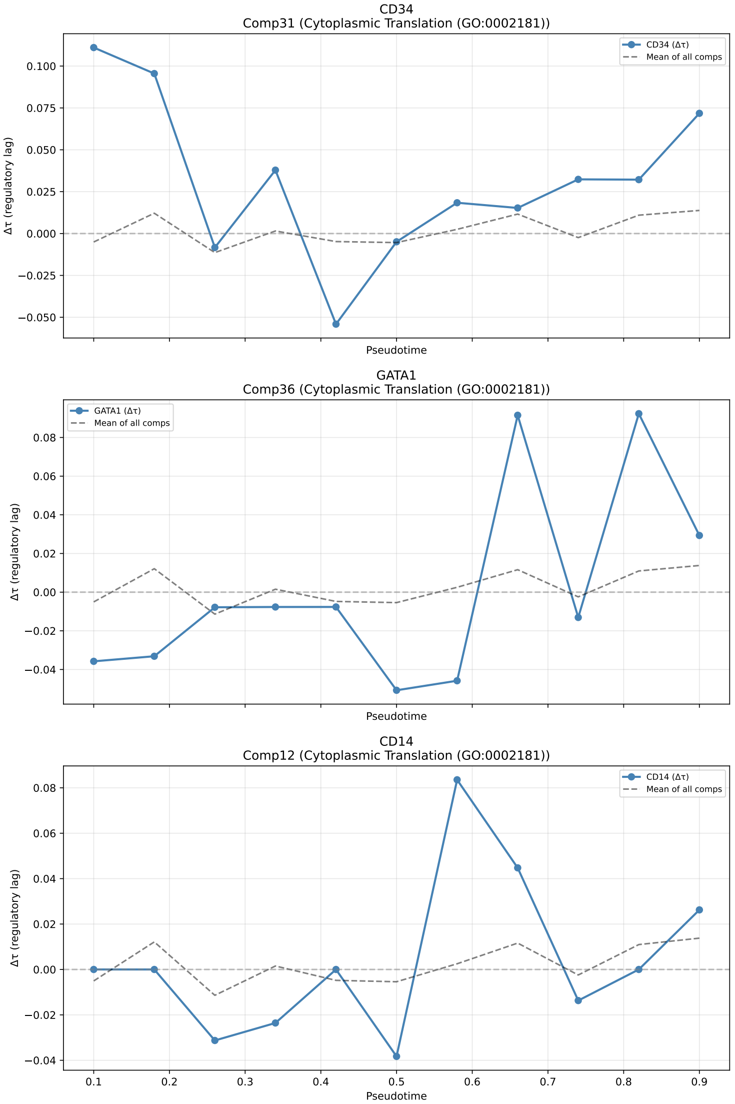


Figure 11: Human PBMC: marker $\Delta\tau$ profiles (CD34, GATA1, CD14).

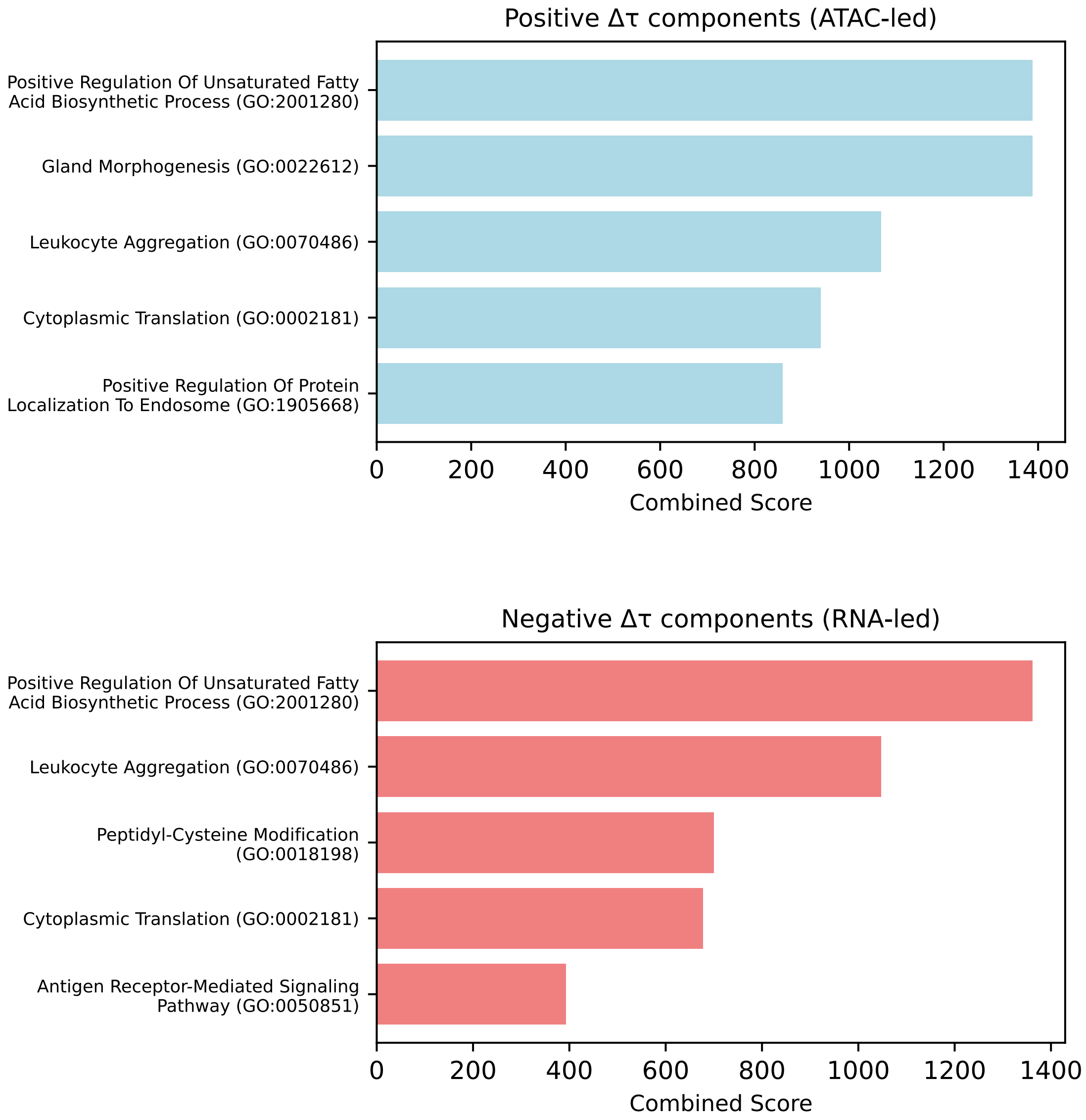


Figure 12: Human PBMC: Gene-program GO enrichment.

#### S4 – Mouse kidney

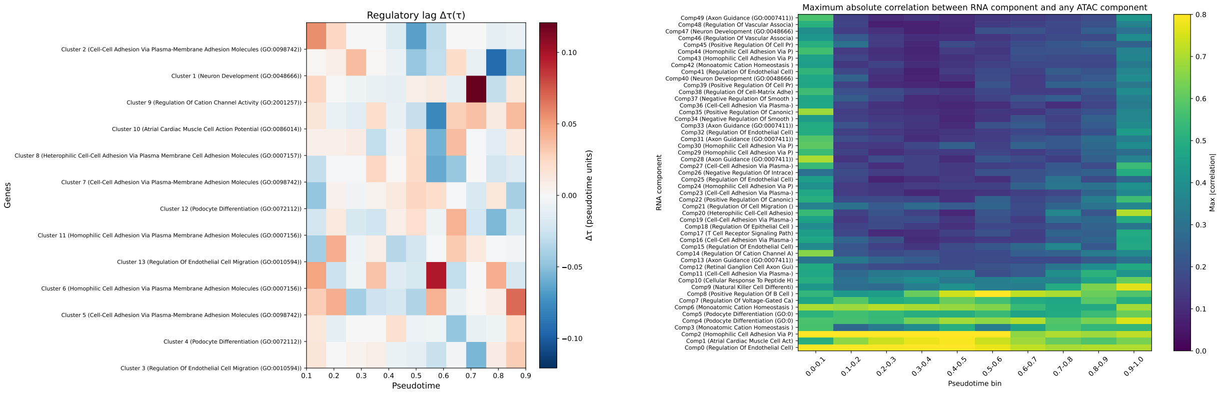


Figure 13: Mouse kidney: $\Delta\tau$ heatmap (left) and ATAC-RNA correlation U-shape (right).

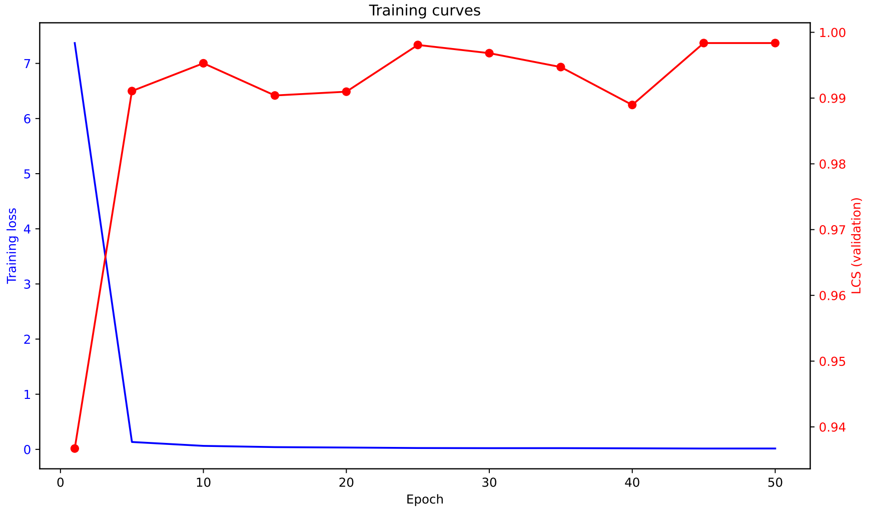


Figure 14: Mouse kidney: training curves (reconstruction loss + LCS).

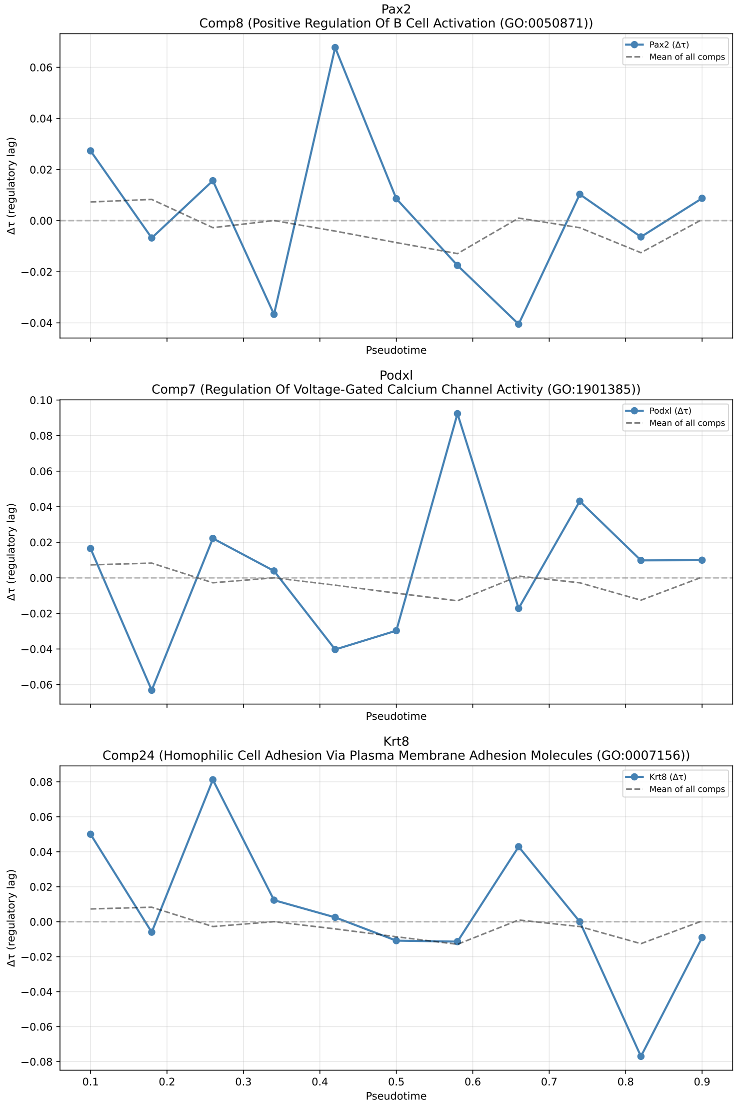


Figure 15: Mouse kidney: marker $\Delta\tau$ profiles (Pax2, Podxl, Krt8).

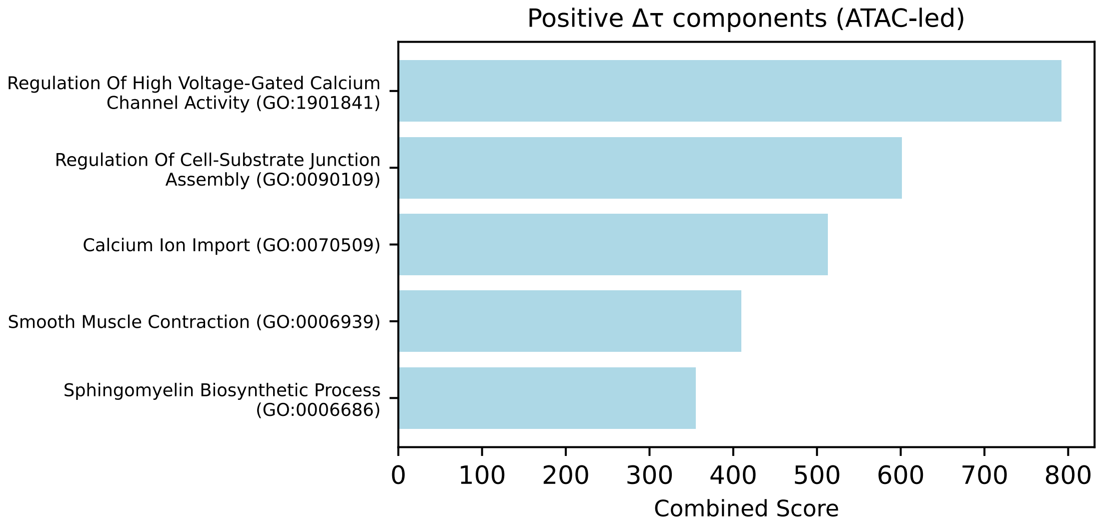


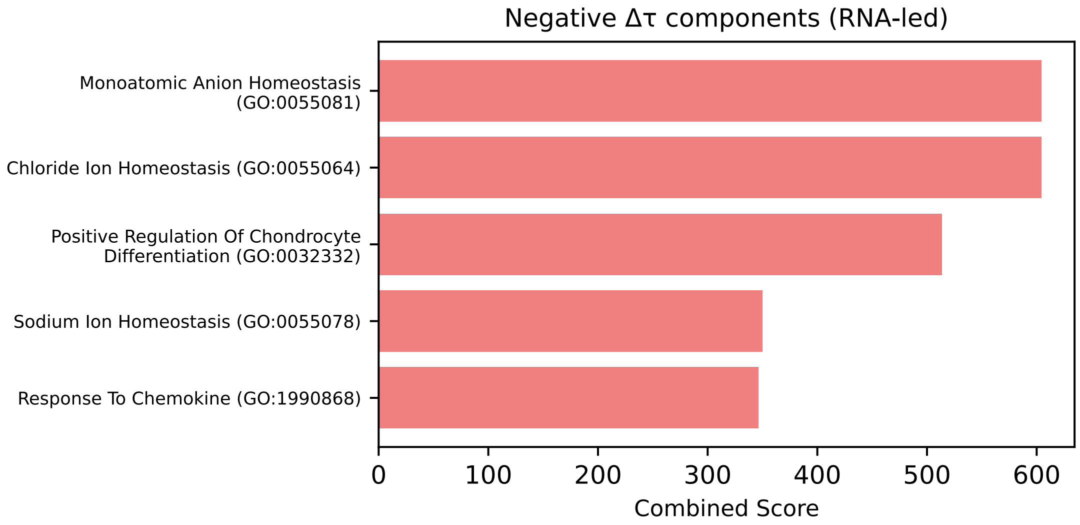


Figure 16: Mouse kidney: Gene-program GO enrichment.

### S5 – LCS scatter plots (predicted vs CCF lags)

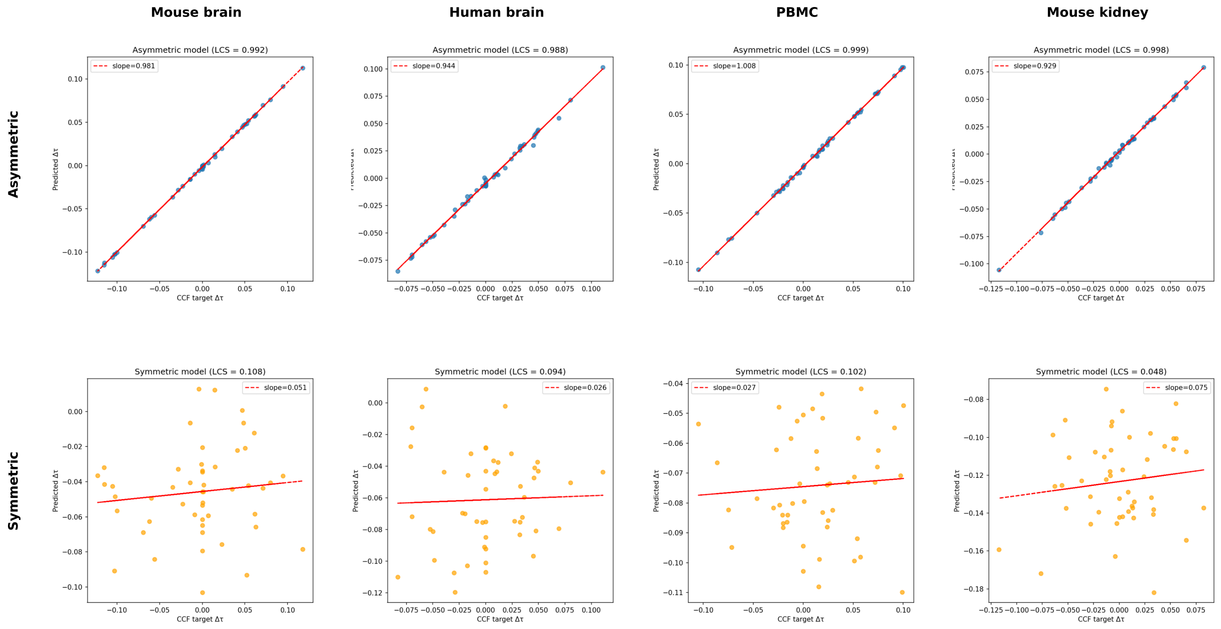


Figure 17: LCS scatter plots for all four datasets. Top row: asymmetric model; bottom row: symmetric model. Red dashed line shows the linear fit; slope and LCS are annotated.

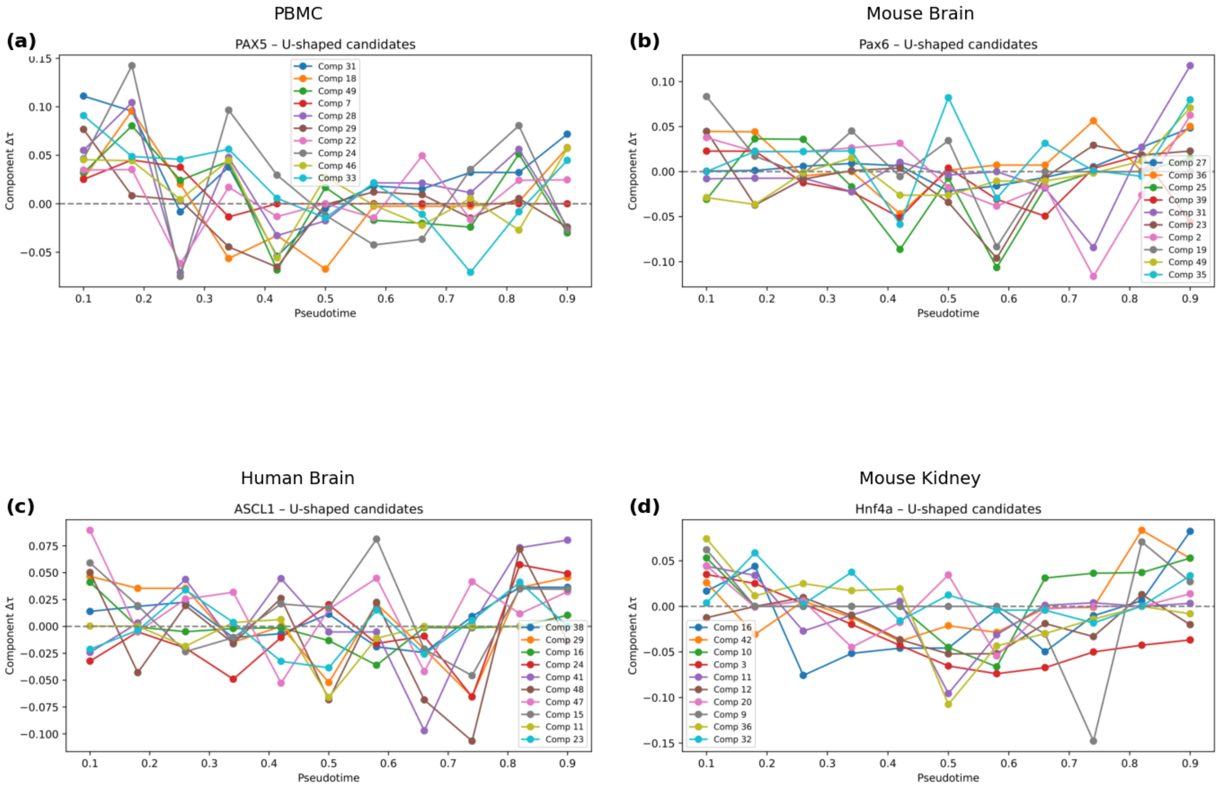


Figure 18: Biphasic regulatory lag profiles for statistically significant components in all four tissues (permutation $p < 0.05$). Each panel shows the component-level $\Delta\tau$ across pseudotime windows; a dashed horizontal line marks the ATAC-led / RNA-led boundary. The corresponding ATAC–RNA correlation U-shapes and permutation p-values are listed in Supplementary Table ST5.

### Supplementary Tables

#### ST1 – Perturb-seq $\delta\Delta\tau$ validation summary

Table 1: Perturb-seq $\delta\Delta\tau$ results (one-sided Mann-Whitney U test).

| Knockout | NTC cells | Perturbed cells | Target $\delta\Delta\tau$ mean | Background $\delta\Delta\tau$ mean | p-value |
|---|---|---|---|---|---|
| SMARCB1 | 1,144 | 147 | $3.0 \times 10^{-4}$ | $2.0 \times 10^{-4}$ | 0.0556 |
| SMARCE1 | 25,125 | 3,394 | $\sim 0$ | $\sim 0$ | 0.0089 |

#### ST2 – Lag Generalization

Table 2: Held-out generalization results (80/20 stratified split).

| Dataset | Train cells | Test cells | Held-out LCS (training target) | Test-set CCF LCS |
|---|---|---|---|---|
| Human PBMC | 2,402 | 606 | 0.9885 | −0.2092 |
| Mouse brain | 3,899 | 981 | 0.9484 | −0.1556 |
| Human brain | 2,661 | 670 | 0.8483 | −0.0252 |
| Mouse kidney | 4,224 | 1,062 | 0.9369 | 0.2214 |

#### ST3 – Full Model vs. Cell-state Ablated Model

Table 3: Cell-state ablation results across all four datasets.

| Dataset | Full LCS | Ablated LCS | Mean \|slope\| full | Mean \|slope\| ablated | KS $p$-value |
|---|---|---|---|---|---|
| Mouse kidney | 0.9984 | 0.7136 | 0.0404 | 0.0092 | $2.0 \times 10^{-27}$ |
| Mouse brain | 0.9920 | 0.8267 | 0.0142 | 0.0060 | $3.2 \times 10^{-19}$ |
| Human brain | 0.9877 | 0.9291 | 0.1114 | 0.0051 | $2.0 \times 10^{-29}$ |
| Human PBMC | 0.9990 | 0.9112 | 0.0026 | 0.0044 | $3.9 \times 10^{-2}$ |

Table 4: ST4 – Gene-level CCF validation results across all four datasets.

| Dataset | TF | Gene-level CCF $p$ | TMO $p$ (component) | Variance ratio (gene/TMO) |
|---|---|---|---|---|
| Human PBMC | PAX5 | $6.3 \times 10^{-40}$ | $3.0 \times 10^{-23}$ | 3.80 |
| Mouse brain | Pax6 | $2.9 \times 10^{-38}$ | $2.4 \times 10^{-18}$ | 4.67 |
| Human brain | ASCL1 | $8.3 \times 10^{-11}$ | $1.0 \times 10^{-3}$ | 4.19 |
| Mouse kidney | Hnf4a | $1.1 \times 10^{-14}$ | $2.0 \times 10^{-4}$ | 3.68 |

Table 5: ST5 – Statistically significant biphasic regulatory lag components.

| Dataset | Component | Template $r$ | Permutation $p$ | Correlation dip (early → mid → late) |
|---|---|---|---|---|
| Human PBMC | 31 | 0.765 | 0.005 | 0.65 → 0.09 → 0.55 |
| Human PBMC | 18 | 0.757 | 0.004 | 0.69 → 0.14 → 0.53 |
| Mouse brain | 27 | 0.603 | 0.026 | 0.75 → 0.13 → 0.55 |
| Mouse brain | 36 | 0.584 | 0.037 | 0.66 → 0.12 → 0.53 |
| Human brain | 38 | 0.778 | 0.002 | 0.48 → 0.10 → 0.36 |
| Human brain | 29 | 0.680 | 0.005 | 0.50 → 0.16 → 0.48 |
| Human brain | 16 | 0.610 | 0.006 | 0.41 → 0.15 → 0.77 |
| Mouse kidney | 16 | 0.802 | 0.001 | 0.47 → 0.11 → 0.48 |
| Mouse kidney | 42 | 0.659 | 0.021 | 0.44 → 0.10 → 0.32 |
| Mouse kidney | 10 | 0.607 | 0.015 | 0.50 → 0.29 → 0.61 |
| Mouse kidney | 3 | 0.598 | 0.026 | 0.57 → 0.27 → 0.53 |
| Mouse kidney | 11 | 0.593 | 0.015 | 0.48 → 0.19 → 0.42 |
| Mouse kidney | 12 | 0.575 | 0.026 | 0.49 → 0.18 → 0.49 |
